# Discovery And Characterization of Small Molecule Inhibitors of Zika Virus Replication

**DOI:** 10.1101/2022.12.15.520558

**Authors:** Ankur Kumar, Deepak Kumar, Prateek Kumar, Brittany L. Jones, Indira U Mysorekar, Rajanish Giri

**Affiliations:** School of Basic Sciences, Indian Institute of Technology Mandi, VPO-Kamand, HP 175005, India; Department of Medicine, Section of Infectious Diseases, Baylor College of Medicine, Houston TX 77030; Department of Molecular Virology and Microbiology, Baylor College of Medicine, Houston TX 77030

**Keywords:** Zika virus, NS3 protease, NS3 helicase, MTase, RdRp, flavivirus, drug targets, small molecule inhibitor

## Abstract

Zika virus (ZIKV) is a flavivirus, and ZIKV infections in the past 15 years have been linked to Guillain-Barre syndrome and severe complications during pregnancy associated with congenital Zika syndrome. There are no approved therapies or vaccines for ZIKV. In recent years, advances in structure-based drug design methodologies have accelerated drug development pipelines for identifying promising inhibitory compounds against viral diseases. Among ZIKV proteins, NS2B-NS3 protease is an attractive target for antiviral drug development due to its vital role in the proteolytic processing of the single polyprotein. To find potential inhibitors against ZIKV, we used molecular docking at the NS2B-NS3 protease active site as a virtual screening approach with small molecules diverse scaffold-based library with rigorous druglikeness filters. The top-hit compounds with stable molecular dynamics trajectories were then subjected to in-vitro efficacy testing against ZIKV. In docking and molecular dynamics simulation studies, compound F1289-0194 showed stable binding to the NS2B-NS3 protease active site. Furthermore, viral load assays, immunofluorescence, and plaque reduction assays demonstrated that compound F1289-0194 significantly reduced ZIKV load and replication in Vero cells while maintaining cellular integrity. Thus, the compound F1289-0194 merits further investigation as a novel inhibitor against ZIKV replication.

## 1. Introduction

Following the first major Zika virus (ZIKV) outbreak in 2007 (1) and the subsequent epidemic of 2015-16, no therapeutics or vaccines have become available to treat or prevent ZIKV infection specifically. Although the number of ZIKV cases has diminished globally, transmission continues in regions where it has become endemic, outbreaks in new regions continue to occur, and significant risks remain for future epidemics (2–4). This is cause for alarm, particularly among pregnant individuals, due to the virus’s association with developmental and neurological abnormalities in the fetus (5–7).

ZIKV is an enveloped positive-strand RNA arbovirus in the Flaviviridae family. ZIKV RNA is synthesized within virus-induced replication vesicles derived from the endoplasmic reticulum (8–10). The viral genome encodes a single polyprotein that is hydrolyzed by the viral and host proteases into three structural (C, prM, E) and seven non-structural (NS1, NS2A, NS2B, NS3, NS4A, NS4B, NS5) proteins (11, 12). Of these, the non-structural protein NS3 is an excellent drug target as it has protease and helicase domains and possesses enzymatic activity for ZIKV replication.

The NS3 protein has protease and helicase domains, thus making it an excellent drug target due to its multifunctional role in viral replication (13, 14). As a protease, NS3 is a key enzyme for the proteolytic processing of the polyprotein, which requires NS2B as a co-factor for its enzymatic activity (15–17). Together, NS2B and NS3 protease form the NS2B-NS3 protease complex. Apart from this proteolytic function, the NS2B-NS3 protease complex is thought to play other functional roles, including involvement in the replication complex formation, interaction with other viral proteins, NS4B and NS5, interaction with the polymerase, and in modulation of host immune response and pathogenesis (18–21).

In addition, NS3 also functions as a helicase, which is required to unwind the RNA at the time of viral RNA synthesis, while the NTPase activity of the helicase domain provides energy for RNA intermediate unwinding (20, 22). The structural characterization of the NS3 helicase suggests that the substrate-binding site, the RNA binding site, and ATP binding site would be important targets for the inhibitor discovery (23). Previously, some small molecules and nucleoside analogs have been identified that have the potential to inhibit the helicase and NTPase activity in Hepatitis C virus and dengue virus (24–27). Therefore, NS3 helicase inhibition could be important in inhibiting viral replication in ZIKV.

NS5 is the largest protein encoded by the ZIKV and has a crucial role in viral replication, making it another potential target for inhibitor discovery. The NS5 protein has two domains, MTase and RdRP, where both domains perform a specific functional role in replication. The MTase domain methylates the RNA cap, enabling the virus to evade the host immune response, and this characteristic makes MTase an attractive drug target (28, 29). The RdRp domain synthesizes the (-) sense RNA from the (+) sense template, and the synthesized strands are used to translate the polyprotein and replicate (+) sense RNA for the packing as viral genome into the nascent virus particle (30, 31). This domain also interacts with NS3 and host proteins and acts as an antagonist for IFN response (22, 32). Diversity in the functional roles the RdRp domain an attractive target for antiviral therapeutics against ZIKV.

In contrast to many prior approaches that have explored the use of repurposed drugs (33), our objective here is to discover novel compounds that could target NS3 or NS5 to inhibit ZIKV replication. To achieve this, we took a computational drug discovery approach using docking and simulation coupled with further *in vitro* validation using virological assays. A key caveat with drug discovery studies is potential cellular toxicity limiting their application. Here, we identify compounds with an anti-protease activity that not only limits ZIKV load but is safe to use in human cells.

## 2. Material and methods

### 2.1. Site mapping and grid generation

For NS5 MTase (PDB ID: 5KQR) and NS5 RdRp (PDB ID: 5WZ3), we have generated a site map using the SiteMap software module (Schrödinger) and based on the site score, druggability score, and key residues in predicted sites, best binding site was chosen. A grid was generated on the selected binding site of both enzyme domains. The grid center coordinates X, Y, and Z for the grid on NS5 MTase corresponded to 13.04, 5.02, and 5.84 Å, and for NS5 RdRp corresponded to 35.71, 3.49, and 76.27 Å, respectively. For NS2B-NS3 protease (PDB ID: 5LC0), we used the same grid (ligand binding site) on the active site of NS2B-NS3 protease, as taken in our previous study (34). For NS3 helicase (PDB ID: 5GJC), we used our previously reported NTPase site where ATP hydrolysis occurs during the helicase enzyme activity (35, 36).

### 2.2. Ligand preparation

We used a library of small molecules (18098 compounds) from the LifeChemicals compound database. The compounds in the LifeChemicals library distinguish themselves from others because they are drug-like small molecules with high binding affinities to their targets, as predicted by ligand-based or receptor-based techniques. They also have compounds that share scaffolds with other reported targeted molecules, such as those having antiviral and anticancer properties. These compounds were prepared using the Epik program (Schrödinger) to calculate protonation states and OPLS 2005 Force Field (Schrödinger) for ligand minimization and stereoisomers.

### 2.3. Molecular docking

The glide scoring function of the Schrödinger suite is highly accurate and reliable for screening molecules and identifying hits based on atomic-level interaction and energy-based scoring before performing an experimental investigation. We used three different precisions of virtual screening sequentially: High Throughput Virtual Screening (HTVS), Standard Precision (SP), and Extra Precision (XP). Initially, all the molecules were docked against the catalytic site of NS2B-NS3 protease enclosed by a grid, as mentioned in section 2.1. Upon docking, the compounds showing the best docking score, binding energy, and interactions with the catalytic residues of ZIKV NS2B-NS3 protease were manually screened. The interaction of the top 10 molecules based on the docking score from each library is summarized in Table S1. Among the 30 top compounds, F1289-0194, F2477-0054, F3260-0474, and F6252-4720 were selected based on their low molecular weight and higher similarities with the existing compounds in the PubChem compound database. These four compounds were docked with NS3 helicase, NS5 MTase, and NS5 RdRp to analyze the binding interaction with these proteins further.

### 2.4. Binding-free energy calculation

Using approaches similar to those used previously by members of our group (37–39), we calculated the binding free energy of ligands binding at the protease active site. The obtained poses were subjected to binding energy calculation using the Prime utility (Schrödinger), which employs the MM-GBSA method to calculate the binding energy of the complex (40). Prime was shown to be a very efficient tool to calculate the binding energy of the protein-ligand complex and provide the best pose of a ligand with the protein.

### 2.5. Molecular dynamics simulation

The Desmond simulation package embedded in the Schrödinger suite was used for MD simulation by following previously described protocols (38). We used NS2B-NS3 protease docked complex with compounds F1289-0194, F2477-0054, F3260-0474, and F6252-4720 for MD simulation. Briefly, the protein-ligand complex was enclosed in the center of an orthorhombic box with an edge distance of 10, and the rest simulation box was filled with a TIP4P water model. The setup included 0.15 M NaCl and charges neutralized with counterions for proper electrostatic distribution. The steepest descent method was employed for energy minimization with up to 5000 iterations containing ten steps. Finally, the production run was carried out for 200 ns using the Nose-Hoover thermostat and Martina-Thomas-Klein (MTK) barostat methods for temperature and pressure coupling throughout the simulations.

### 2.6. NS2B-NS3 protease inhibition assay

NS2B-NS3 protease inhibition assay (see the purification protocol as described previously (41)) was performed in 10 mM Na-phosphate buffer (pH 7) containing 1 mM TCEP, 1 mM CHAPS, and 20 % glycerol. NS2B-NS3 protease was incubated for 20 minutes with the respective compounds. Afterward, the compound-NS2B-NS3 protease complex was mixed with the substrate (Bz-nKKR-AMC; for details, see the previous report (34)). In 50 μL reaction volume, the final NS2B-NS3 protease concentration was kept at 5nM, and the reaction was performed in triplicates in a multi-well plate (complete black 96 well). The fluorescence intensity of the reaction was monitored using a multi-well plate reader (infiniteM200PRO: TECAN) with an excitation wavelength of 360 nm and emission wavelength of 460 nm at every 5-minute interval. Fluorescence (F_t_ - F_o_; F_o_: initial fluorescence, F_t_: Fluorescence after 15 minutes) was converted to nM of AMC release using a standard curve plotted using fluorescence of free AMC. Further, velocity (AMC release [nM]/minute] was plotted as the function of substrate [μM] to compute inhibition constant (*K_i_*) using GraphPad Prism.

### 2.7. Cell viability assay

Vero cells were grown in DMEM/F12 culture media supplemented with 10% FBS, incubated in a 5% CO_2_ incubator at 37 °C, and passaged at 80-90% confluency. Cells were seeded in a flat bottom 96 well plate (100 μL/well culture media volume) at 4,000 cells/well and incubated overnight in a 5% CO_2_ incubator at 37 °C. On the next day, 100 μL of two-fold serially diluted compound samples in DMEM/F12 media were added to the respective well at a final concentration range of 3.25 to 200 μM and incubated further for 48 hrs. To each well, 15 μL of the 12 mM MTT stock solution was added, and cells were incubated at 37 °C. After 4 hours, media was removed from the wells, and the formazan was dissolved in 100 μL of 100% DMSO. Afterward, the absorbance was monitored 10 minutes postincubation at 540 nm, and a reference was taken at 640 nm. The following equation used the subtracted absorbance (Abs540-Abs640) to estimate the percentage of cell viability.

% Cell viability = [ Abs _treated_/ Abs _non-treated_] * 100

### 2.8. ZIKV infection

As used in our previous report, a Brazilian strain (Paraiba 2015) of the ZIKV was used to infect the Vero cells under biosafety level 2 (34). Incubation at each step was done at 37 °C in a 5% CO_2_ incubator. Wells in a 24-well plate were seeded with 40,000 Vero cells/well using DMEM/F12 culture media supplemented with 10% FBS and incubated overnight. Infection was done at 0.1 multiplicity of infection (MOI) with 0.5 ml of culture media without FBS, incubated for one hour, and then washed twice with PBS. Cells were treated with fresh media containing 12.5 to 100 μM of the tested compounds. Dilutions of the compounds were prepared from 50 mM stock (in 100 % DMSO) by two-fold serial dilution in DMEM/F12 media supplemented with 10% FBS. The supernatant was collected 48 hours post-infection (hpi) for RNA extraction and stored at −80 °C until further use.

### 2.9. Estimation of viral load

Viral RNA was isolated from 140 μL of supernatant from the 48 hpi sample using the QIAamp Viral RNA Mini Kit (Qiagen), following the included protocol. RNA levels were quantified by one-step qRT-PCR on an ABI 7500 Fast Instrument (40 cycles under standard cycling conditions) using TaqManRT enzyme Mix with primer/probe for the ZIKV as described previously (34). The viral load (equivalent per ml) in the supernatant was determined by a standard curve produced by the 10-fold serially diluted ZIKV RNA from known stock and expressed as a log10 scale. The data were analyzed using GraphPad Prism by one-way ANOVA Dunnett’s test. Data were statistically significant for the p-value < 0.05.

### 2.10. Immunofluorescence

Vero cells were grown on chambered slides (Millipore Sigma) for the immunofluorescence studies. ZIKV infection and compound treatment were similar to the experimental setup in 24-well plates. Treated cells, 48 hpi were fixed with 4% paraformaldehyde after washing three times with PBS. Cells were blocked with 1% BSA for 1 hour and incubated overnight at 4°C with the primary antibody for the capsid protein of ZIKV, followed by a secondary antibody labeled with Alexa 488 fluorophore for 1 hour at room temperature. Cells were stained with DAPI and mounted with mounting media. The fluorescent image was taken using a Zeiss LSM880 Confocal Laser Scanning Microscope. The data were analyzed using GraphPad Prism by one-way ANOVA Dunnett’s test. Data were considered to be statistically significant for the p-value < 0.05.

### 2.11. Plaque assay

In a 6-well plate, 5 × 10^5^ cells/well were seeded and grown overnight. Afterward, the media was removed and washed with PBS. The supernatant sample of ZIKV-infected cells with different compound concentrations was taken and serially diluted 10-fold in DMEM/F12 culture media. Then infection was done with 0.5 ml of serially diluted ZIKV sample per well and incubated for 1 hr. The media was removed, cells were washed twice with PBS, overlaid with 4 ml of overlay media (1:1 ratio of mixed 2% Oxoid agar and 2X MEM media, at 54°C), and incubated for 3-4 days at 37°C with 5% CO_2_. Cells were fixed with 1 % formaldehyde solution (1-2 ml) for 1 hour. The formaldehyde was discarded, and the overlay plug was carefully removed using a spatula. To each well, 1 ml of crystal violet solution was added, and incubation proceeded for 10 minutes on the rocker. Excess crystal violet solution was discarded and rinsed in each well with distilled water. Plaques were counted and estimated as a plaque-forming unit (PFU)/ml.

## 3. Results

### 3.1. Virtual screening of small molecules

For virtual screening, we initially used NS2B-NS3 protease as a target for docking the ligand at the binding site where a boronate inhibitor was bound in its crystal structure (PDB ID: 5LC0). Based on the docking score, binding energy, interacting amino acid residues, similar molecules in the PubChem database, and molecular weight we selected F1289-0194, F2477-0054, F3260-0474, and F6252-4720 as potential candidates (Figure S1). These compounds are stabilized at the active site of NS2B-NS3 protease by forming H-bonds, salt bridges, π-π bonds, and hydrophobic interactions with crucial amino acid residues (Figures 1 & S2). The docking score of the selected compounds, F1289-0194, F2477-0054, F3260-0474, and F6252-4720, is found to be −6.98, −6.95, −6.85 and −6.01 kcal/mol, respectively. Importantly, these compounds also interact with the catalytic residues (Asp75, His51) of NS3 protease. The interaction of the key amino residues of the NS2B-NS3 protease with these compounds is summarized in Table S1. These small molecules show stable binding at the NS2B-NS3 protease active site determined by molecular docking and simulation. We further checked the likelihood of the interaction of these molecules with other ZIKV enzymes. So, using the same molecular docking strategy, we docked all four compounds onto the druggable sites of the ZIKV NS5 MTase, NS5 RdRp, and NS3 helicase. Here, F1289-0194 showed good binding affinity at the druggable sites of these enzymes with multiple interacting bonds (Figure 2). The H-bond, salt bridge, π-π bonds, π-cation bonds, and hydrophobic interaction with the amino acid residues of these enzymes stabilize all these compounds. Moreover, F1289-0194 has shown a higher docking score of −7.03 kcal/mol against MTase, indicating that in addition to NS2B-NS3 protease, the NS5 MTase of ZIKV could also be a target for this compound. Compound F2477-0054 has shown a docking score of −5.66 kcal/mol, −4.34 kcal/mol, and −4.25 kcal/mol at the MTase, RdRp, and helicase, respectively (Figure S3). F3260-0474 has shown a docking score of −4.58 kcal/mol, −4.15 kcal/mol, and −3.17 kcal/mol at the MTase, RdRp, and helicase, respectively. F6252-4720 has shown a docking score of −4.28 kcal/mol, −2.85 kcal/mol, and −4.11 kcal/mol at the MTase, RdRp, and helicase, respectively (Figure S3).

**Figure 1.**
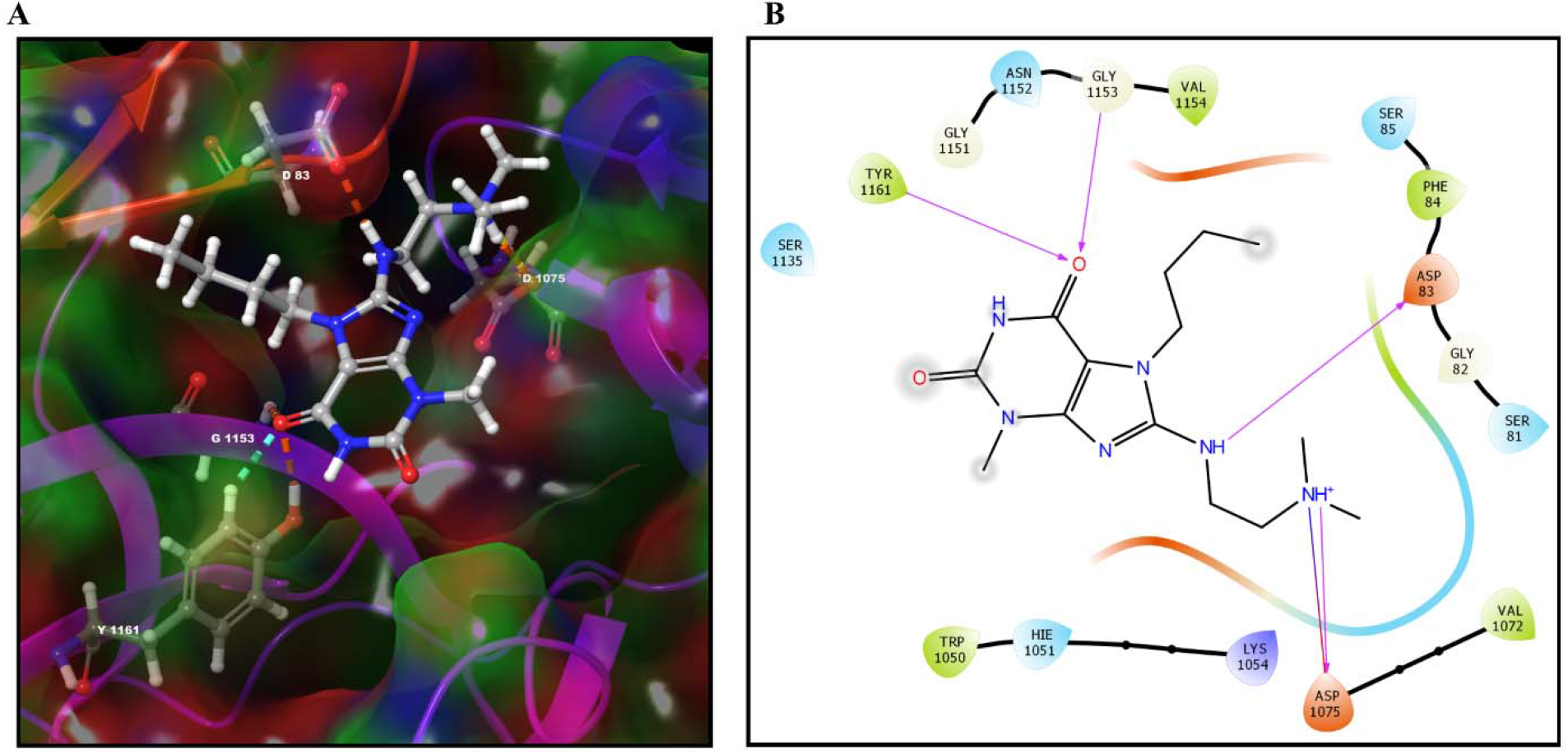
Molecular interaction of F1289-0194 at the active site of NS2B-NS3 protease of ZIKV. (A) 3D interaction diagram of NS2B-NS3 protease highlighting residual contacts via dashed lines. (B) 2D molecular interaction diagram of NS2B-NS3 protease complexed with F1289-0194. The F1289-0194 compound at the center is stabilized by the following interactions with amino acids of the NS2B-NS3 protease: H-bonds (magenta arrow; Asp75, Tyr161, and Gly153 of NS3 protease, and Asp83 of NS2B), one salt bridge (red-blue solid line; Asp75 of NS3 protease), and hydrophobic interactions (Trp50, Val72, Tyr161, and Val154 of NS3, and Phe84 of NS2B). NS2B residues correspond to 49-87; NS3 protease residues are denoted as 1015-1167 and correspond to 15-167.

**Figure 2.**
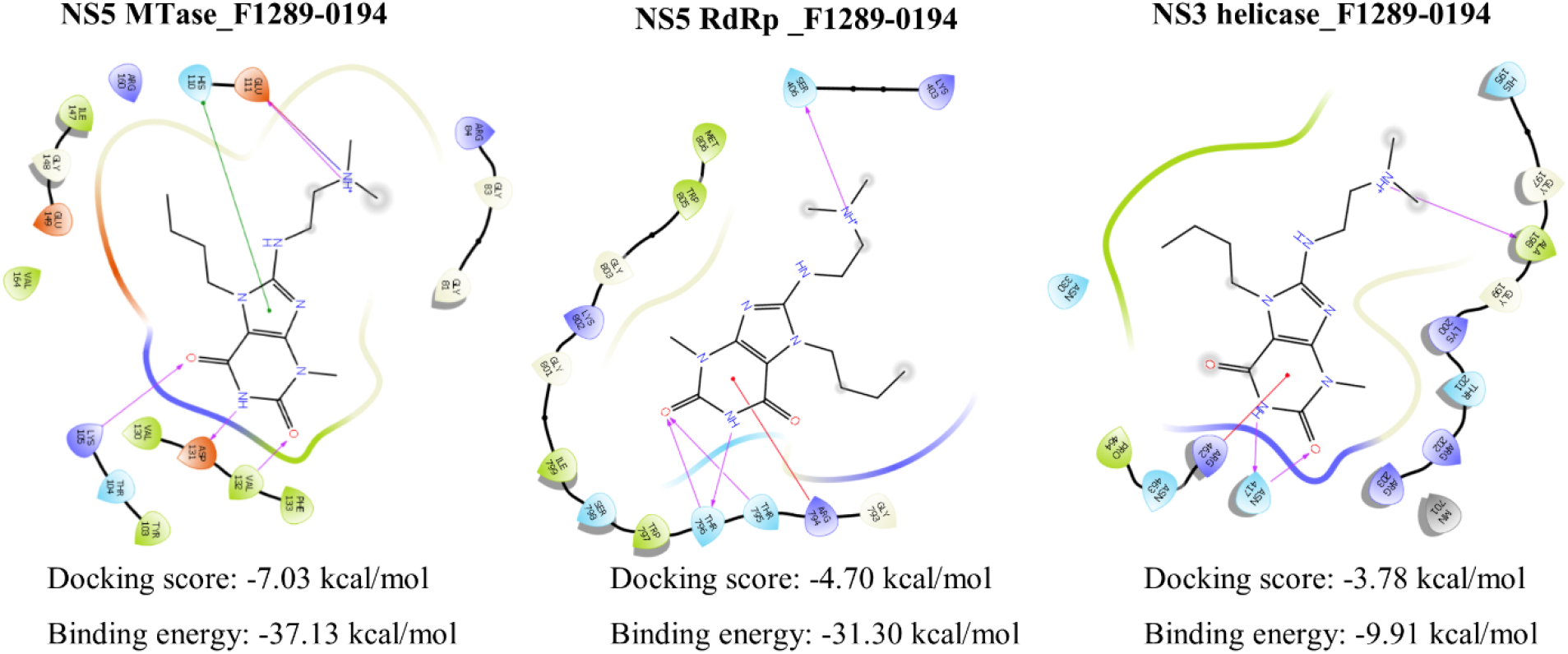
Molecular interaction of F1289-0194 with NS5 MTase, NS5 RdRp, and NS3 helicase of ZIKV. The F1289-0194 compound at the center interacts with the amino acids of these enzymes by H-bond (magenta arrow), salt bridge (red-blue solid line), π-π bond (solid green line), π-cation (solid red line), and hydrophobic interactions. F1289-0194 is showing interaction with NS5 MTase by four H-bonds (lys105, Asp131, Val132, and Glu111), one salt bridge (Glu111), and one π-π bond (His110), with NS5 RdRp by four H-bonds (Ser406, Thr795, and Thr796), and one π-cation bond (Arg794), and NS3 helicase by three H-bonds (Asn417, and Ala198), and one π-cation bond (Arg462). (PDB ID: 5KQR for NS5 MTase; 5WZ3 for NS5 RdRp; 5GJC for NS3 helicase).

### 3.2. Molecular dynamics simulation shows stable binding of selected small molecules at NS2B-NS3 protease active site

The stability of ligand-protein interactions was further analyzed by Molecular dynamics (MD) simulation over 200 ns (Figure 3 & Figure S4, S5). All four selected compounds complexed with NS2B-NS3 protease were subjected to simulation to examine the dynamic behavior and conformational change. The RMSD plot revealed that compounds F1289-0194, F2477-0054, and F3260-0474 demonstrated stability at the active site of NS2B-NS3 protease with deviation from ~1.5 to 2.0 Å (Figure 3A & S4A). F6252-4720 showed low stability at the active site with an RMSD of ~4 Å (Figure S4A). Except for F6252-4720, all the compounds showed minimal fluctuating RMSF values (Figure 3B & S4B). Rg revealed that the compactness of the ligand-protein complex for all the compounds was maintained over the simulation. F6252-4720 showed high Rg at time points between ~110-120 ns but later decreased drastically and maintained compactness again (Figure 3C & S4C). Ligand-protein complex formed 2 to 4 H-bonds in docking (Figures 1 & S2), and these bonds with NS2B-NS3 protease were maintained during simulation (Figures 3D & S4D). In addition, these molecules also maintain their interaction like hydrophobic, ionic, and water bridge contacts with the NS2B-NS33 protease active site over the simulation (Figure S5). F1289-0194 maintain its contact predominantly with Trp 50, His51, lys54, Tyr68, Trp69, Gly70, Val72, and Asp 75 of NS3 protease during the course of simulation (Figure 3D). F2477-0054 maintain its contact predominantly with Glu43, Cys80, Lys84, Asp129, Tyr130, Gly133, Thr134, Ser135, Tyr150, Gly151, Val155 and Tyr161 of NS3 protease during the course of simulation (Figure S5B). F3260-0474 maintain its contact predominantly with Asp83 and Phe84 of NS2B cofactor and His51, Asp75, Asp129, Ser135, Gly151, Asn152, Gly153, Val154, Val155, Lys157, Asn158, Gly159, Ser160 and Tyr161 of NS3 protease during the course of simulation (Figure S5C). F6252-4720 maintain its contact predominantly with Val49 and Asp83 of NS2B cofactor and Thr27, Arg28, Gly29, Leu30, His51, Ser56, Ala57, Arg64, Tyr130, Pro131, Ala132, Gly133, Thr134, Ser135, Tyr150, Gly151, Gly153, Val155, Asn158, and Tyr161 of NS3 protease during the course of simulation (Figure S5D). This compound maintains maximum interaction fraction by water bridges, thus making the least stable molecule at the protease active site. Overall, the interactions fraction is higher for F1289-0194 and F2477-0054 among all four compounds.

**Figure 3.**
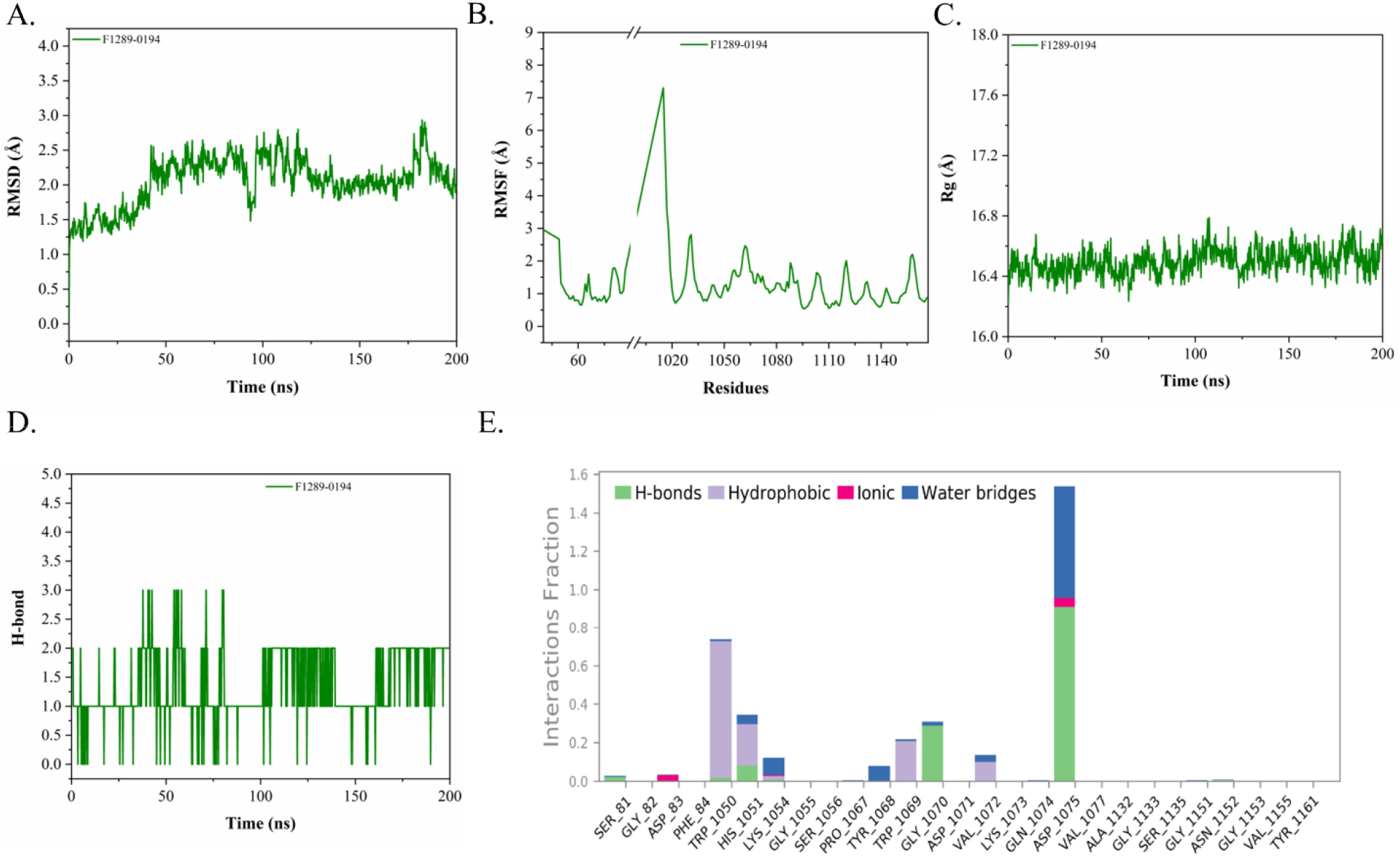
Molecular dynamics simulation of NS2B-NS3 protease complexed with F1289-0194. (A) RMSD represents the stability of the compounds at the active site. (B) RMSF of amino acid residue upon addition of compounds. (C) Rg represents the compaction of NS2B-NS3 protease complexed with the compounds. (D) represents the loss and gain of H-bond during simulation. (E) A bar chart of the interaction of F1289-0194 with the NS2B-NS3 protease residues throughout the simulation of 200 ns. The interactions like H-bonds, Hydrophobic, ionic, and water bridges are maintained with a different fraction of simulation time. For example, a value of interactions fraction of 1 suggests that the contact is maintained 100 % of the simulation time between F1289-0194 and NS2B-NS3 protease active site. If the value is more than 1, it indicates that the same residue forms multiple interactions. [ns: nanoseconds; in panels B & E, NS2B residues correspond to 49-87; NS3 protease residues are denoted as 1015-1167, corresponding to 15-167].

### 3.3. NS2B-NS3 protease inhibition by selected small molecule

The inhibition potential of all four selected compounds against NS2B-NS3 protease of ZIKV was assessed using an enzyme inhibition assay. Enzyme kinetic parameters such as V_max_ and K_m_ for NS2B-NS3 protease are changing in the presence of increasing compound concentration compared to the absence of compounds (Figure S6). To check the mode of enzyme inhibition, we plotted the data points indicated in Figure S6 to the Lineweaver-Burk plot (Figure S7). We found that the compound F6252-4720 shows an inhibition constant (K_i_) of 196.5 ± 29.42 μM for a mixed model of enzyme inhibition. The inhibition constant (K_i_) of the other three compounds (F1289-0194, F2477-0054, and F3260-0474) was not determined may be due to the requirement of very high compound concentration to inhibit protease activity. So, the possibility of the other target, such as MTase, helicase, and RdRp enzyme of ZIKV, must be explored.

### 3.4. Small molecule F1289-0194 inhibits ZIKV replication

In the virtual screening, the compounds F1289-0194, F2477-0054, and F3260-0474 showed a good binding affinity and stability at the protease active site. Based on the highest docking score and stability at the NS2B-NS3 protease active site, we selected compound F1289-0194 for testing the inhibition potential of ZIKV *in vitro*. We checked its toxicity to Vero cells, finding that the cells showed 90-100% viability up to 100 μM of the compound (Figure S8). Thus, we chose a 0 to 100 μM treatment range in the viral assays. Treatment with F1289-0194 at the selected concentration range to ZIKV-infected Vero cells (48 hpi) led to a significant reduction (\*\**p* < 0.01 *at* 12.5 μM, \*\*\**p* < 0.001 *at* 25 μM and \*\**p* < 0.01 *at* 50 μM) in viral load in the supernatant, compared to non-treated (0 μM) sample, shown in Figure 4A. To validate the viral load assay, we examined the inhibition potential of F1289-0194 using immunofluorescence and plaque reduction assays. The cells were imaged, and ZIKV-capsid positive cells were quantified upon treatment with F1289-0194 (25 μM and 50 μM) for 48 hpi. Immunofluorescence analysis showed a significant reduction (\*\*\*\**p* < 0.0001) in ZIKV-capsid positive cells at both 25 and 50 μM treatment with compound F1289-0194 compared to a DMSO control (Figure 4B & C). Also, the plaque reduction assay revealed that the F1289-0194 treatment significantly reduced the number of plaques compared to DMSO and ZIKV controls (Figure 4D) at 10^5^-fold dilution of ZIKV stock. Plaques in each treatment were estimated in PFU/ml (Figure 4E) relative to the non-treated (DMSO/ZIKV only). Compound F1289-0194 did not show its inhibition effect when the Vero cells were treated with the compound before ZIKV infection (Figure 5). This also implies that the compound F1289-0194 does not obstruct ZIKV entry into Vero cells. However, it may interfere with some crucial replication or maturation step in the ZIKV life cycle. From this, we suggest that inhibition of ZIKV replication in Vero cells might be due to broad inhibition of NS2B-NS3 protease and other non-structural proteins of ZIKV.

**Figure 4.**
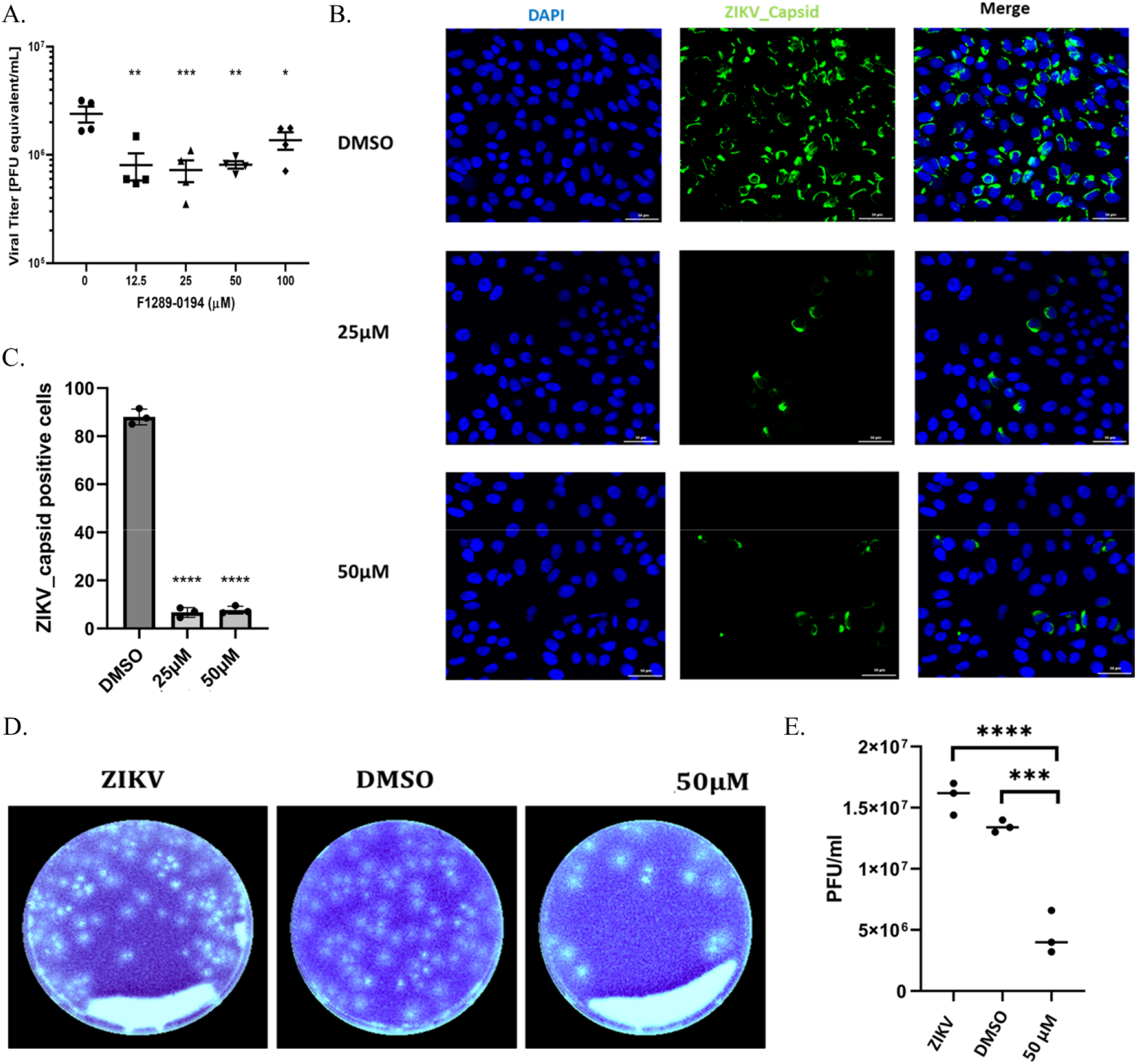
Treatment of F1289-0194 reduces ZIKV replication in Vero cells. (A) The viral burden in the ZIKV-infected Vero cells after treatment with the compounds F1289-0194 for 48 hpi. RNA from the supernatant of indicated treatment of ZIKV-infected cells was quantified by one-step q-RT-PCR and represented as PFU equivalent/ml. Symbols represent each biological replicate, and the bar represents the mean ± standard error of the mean. ns: non-significant; asterisks (*) represent the level of significant difference compared to non-treated (0 μM). \*\**P* < 0.01, \*\*\**P* < 0.001, \*\*\*\**P* < 0.0001 (one-way ANOVA Dunnett’s test). (B) Imaging of ZIKV-infected cells at the indicated treatment of F1289-0194 for 48 hpi. Nuclei are stained in blue, and ZIKV-capsid-positive cells are stained in green. (C) Quantification of ZIKV-capsid positive cells at the indicated treatment. (D) Plaque reduction assay, where the white spot on each plate represents the ZIKV plaque in the respective treatment. (E) Estimation of PFU/ml in the respective treatment of panel D. Bar represents the mean ± standard error of the mean for three biological replicates. Asterisks (*) represent the level of significant difference compared to nontreated (0 μM). \*\*\**P* < 0.001, \*\*\*\**P* < 0.0001 (one-way ANOVA Dunnett’s test).

**Figure 5.**
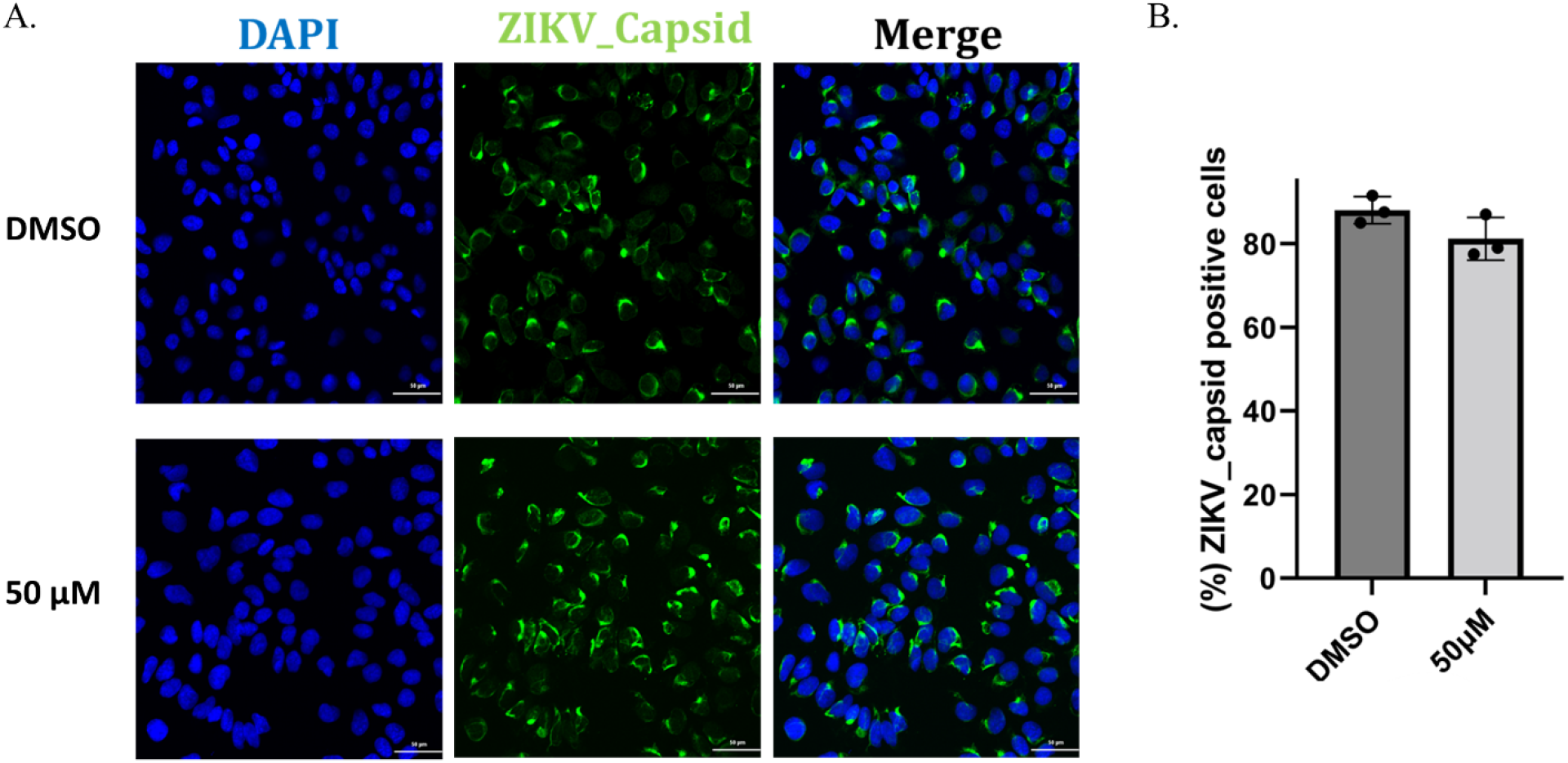
F1289-0194 treatment does not interfere with ZIKV entry to Vero cells. (A) Imaging of ZIKV-infected cells at the indicated treatment before ZIKV infection. Nuclei are stained with blue, and ZIKV-capsid positive cells are stained green. (B) Quantification of ZIKV-capsid positive cells at the indicated treatment. The bar represents the mean ± standard error of the mean for three biological replicates.

### 3.5. Inhibition potential of other selected small molecules against ZIKV

We also investigated the inhibition potential *in vitro* for the other three compounds (F2477-0054, F3260-0474, and F6252-4720) from the selected list of the screened compounds against ZIKV. First, we checked their toxicity to Vero cells, where cells showed 90-100 % viability at up to 100 μM of the compound (Figure S8). Thus, we choose a 0 to 100 μM treatment range for these compounds for ZIKV-infected cells in the viral assays. Treating ZIKV-infected Vero cells with these compounds for 48 hpi at the indicated concentration led to a significant (*\*\*p < 0.01*) reduction of viral load in the supernatant compared to nontreated (0 μM) cells (Figure S9).

## 4. Discussion

In the past, many infectious viruses have emerged and re-emerged, with some becoming a global threat to public health (42, 43) e.g., the ongoing pandemic of COVID-19 (44, 45). At the same time, other infectious viruses exist for long periods. For instance, the spread of flaviviruses such as dengue virus, West Nile virus, yellow fever virus, Japanese encephalitis virus, ZIKV, and others have created alarming health situations in the past and continue to emerge in the wild, causing serious threats to the human population (46). The previous successive outbreaks of ZIKV and its association with neurological disorders demand fast-track discovery of specific inhibitors or the development a vaccine candidate. Until now, only a few vaccine candidates have advanced beyond the early phases of clinical trials (33) and remain on the verge of authorization, hampered by questions about efficacy due to antibody-dependent enhancement of the disease severity and waning epidemic (47, 48). These vaccine candidates will also face the added challenge of being clinically evaluated for their safety and efficacy in pregnant women. Therefore, discovering antiviral molecules becomes more critical for treating ZIKV-infected patients.

Identifying a specific target is vital in screening a lead compound that may function as an inhibitor. Many inhibitor candidates have shown the ability to target viral proteins, including the envelope, protease, helicase, and polymerase (33). Among them, viral protease seems to be an attractive target due to its critical role in the proteolytic cleavage of the polyprotein, inhibiting the further onset of the viral replication life cycle. Thus, we initially screened potential small molecules by targeting the ZIKV NS2B-NS3 protease. First, we utilized a structure-based drug discovery approach to screen potential candidates. Then, we shortlisted four leading compounds (F1289-0194, F2477-0054, F3260-0474, and F6252-4720) that have shown stable interaction at the active site of NS2B-NS3 protease. We also analyzed the possible interaction of these four molecules with three other ZIKV targets (NS5 MTase, NS5 RdRp, and NS3 helicase) are also likely to bind with these targets. These compounds are stabilized at NS3 protease active site by non-covalent and hydrophobic interactions with amino acid residues of both NS3 protease and NS2B co-factor. Among the interacting residues, His51, Asp75, and Ser135 residues of NS3 protease are known to form a catalytic triad (49) and were able to interact with these compounds by H-bonds, suggesting that these interactions may be effective in blocking the protease active site and thus may inhibit its activity in-vitro.

Moreover, these compounds also interact with the Asp83 residue of the NS2B co-factor, which is crucial because a point mutation at this site decreases the protease activity, as reported in a previous study(49), indicating that interaction at this residue may have an advantage in inhibiting the protease activity. We have previously reported a similar interaction in which hydroxychloroquine inhibits protease activity (34). Interestingly, the catalytic triad (His51, Asp75, and Ser135) at NS3 protease is required for its activity and has been reported to be conserved among flaviviruses (49, 50). So, blocking the interaction of this triad may provide an additional advantage of targeting the protease for other flavivirus members. In this context, many inhibitors have also been identified that can potentially block NS2B-NS3 protease activity and, in turn, have the potential to inhibit ZIKV replication (34, 51–53). In addition, other drugs have also shown potential as ZIKV inhibitors by different means(33).

To elucidate the specific mechanistic role of the selected four compounds, further experimental validation is needed *in vitro*. Based on a previous report of FDA-approved drug library screening that targets NS2B-NS3 protease (34) and previously reported literature (33) on inhibition of ZIKV replication, we choose to test the antiviral properties of these compounds in ZIKV-infected cells. All four compounds were proven to lower viral load. In addition, we further found that the compound F1289-0194 does not interfere with viral entry but can interfere with the post-entry steps. However, this compound shows good binding affinity to the non-structural proteins like NS2B-NS3 protease, MTase, and RdRp, indicating the inhibition of the post-entry replication step of the Zika virus life cycle.

Many previous pieces of literature warrant investigation of inhibitors of ZIKV that have the potential to target both entry and post-entry steps (33). Such inhibitors have been reported to inhibit viral entry to the host cell by targeting receptor binding, fusion, and endocytosis steps (33, 54–58). Further testing of these compounds is needed to investigate the detailed insight into the mechanism of action and its inhibition potential using *in vivo* model systems. The inhibitors identified in this study may act as starting candidates for drug discovery research to optimize the strategies for treatment and preventive care for ZIKV patients and could be used as a potential candidate to target protease activity in other flaviviruses.

## Abbreviations

ZIKV: Zika virus
MD simulation: Molecular dynamics simulation
RMSD: root mean square deviation
RMSF: root mean square fluctuation
Rg: radius of gyration
PCR: polymerase chain reaction
RdRp: RNA dependent RNA polymerase
MTase: methyltransferase
HTVS: High Throughput Virtual Screening
SP: Standard Precision
XP: Extra Precision
MM-GBSA: Molecular-Mechanics/ Generalized Born Surface Area
PBS: phosphate buffer saline
FBS: fetal bovine serum
PFU: plaque-forming unit
MOI: multiplicity of infection

## Acknowledgments

This work was supported in part by a Fulbright-Nehru Doctoral Research Fellowship to AK; MHRD-SPARC (SPARC/2018-2019/P37/SL) to RG and IUM; NIH/NICHD grant R01 HD091218 and 3R01HD091218-04S1(RADx-UP Supplement) (To IUM); Indian Council of Medical Research (52/04/2020/BIO/BMS) to RG. RG is also grateful Science and Engineering Research Board (SERB) grant, India (CRG/2019/005603); IYBA Award (BT/11/IYBA/2018/06) grant, Department of Biotechnology (DBT), India; Indian Council of Medical Research (58/6/2020/PHA/BMS). Finally, we thank Dr. Robert Lawrence for his valuable comments and editing.

## Conflict of interest

IUM serves on the Scientific Advisory Board of Luca Biologics. No other conflict of interest exists.

## Author Contributions

AK, DK, IUM, and RG designed experiments, and AK and DK performed the majority of experiments, helped by BJ. AK and PK performed virtual screening and simulation of compounds. AK, DK, and PK wrote the first draft with input from RG and IUM. All authors approve of the final draft.

