## Supplemental file for "Discovery And Characterization of Small Molecule Inhibitors of Zika Virus Replication"

**Supporting information**

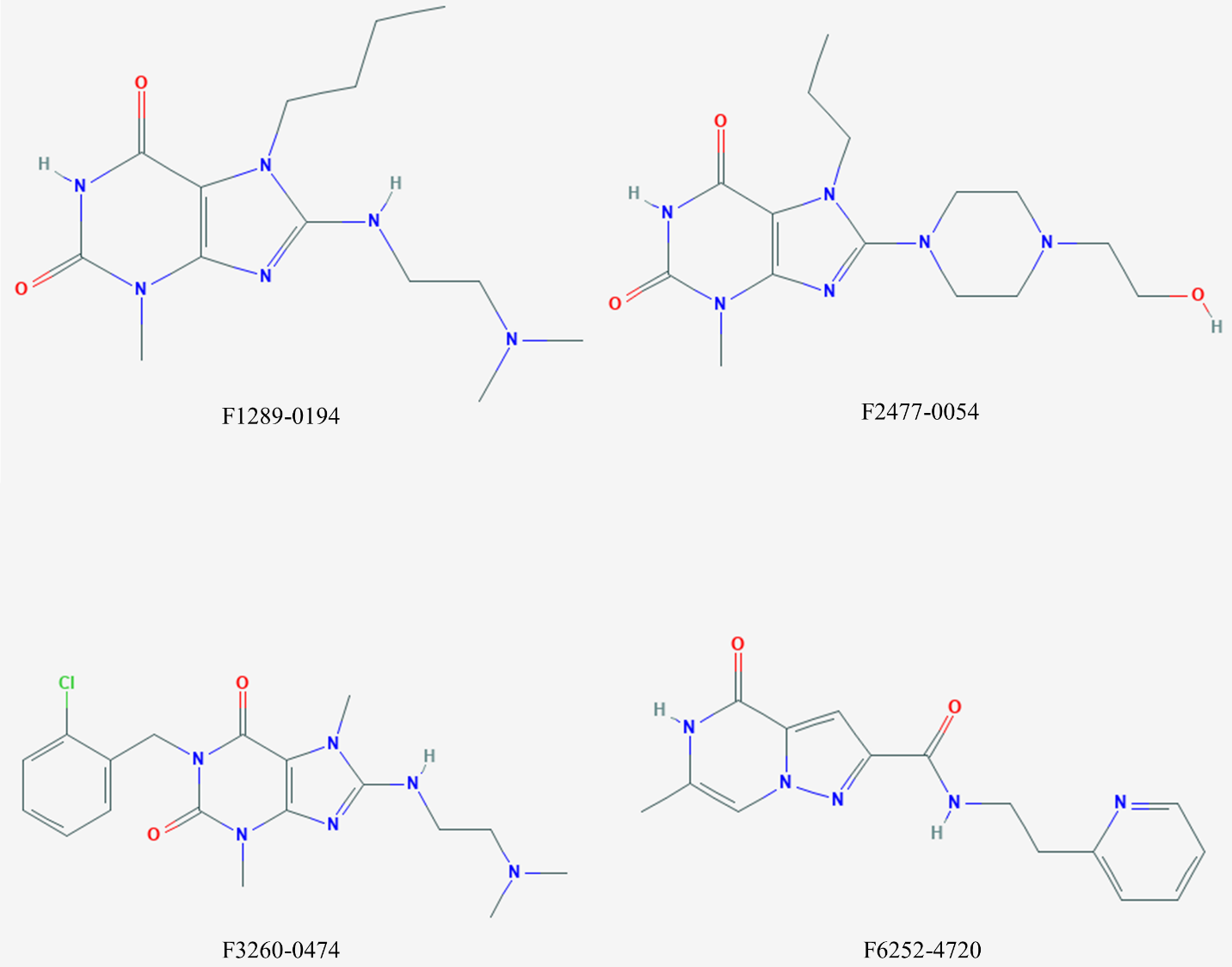

**Figure S1. Shortlisted compounds from virtual screening.** The structure of all compounds is taken from the PubChem database.

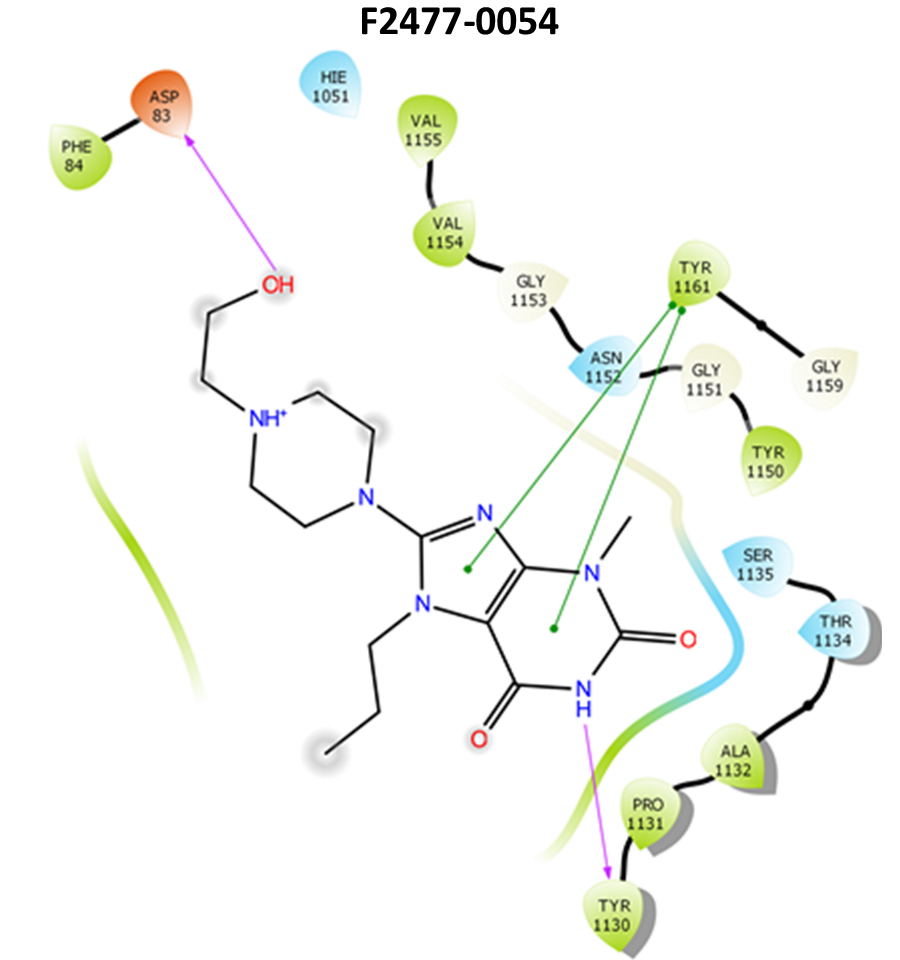

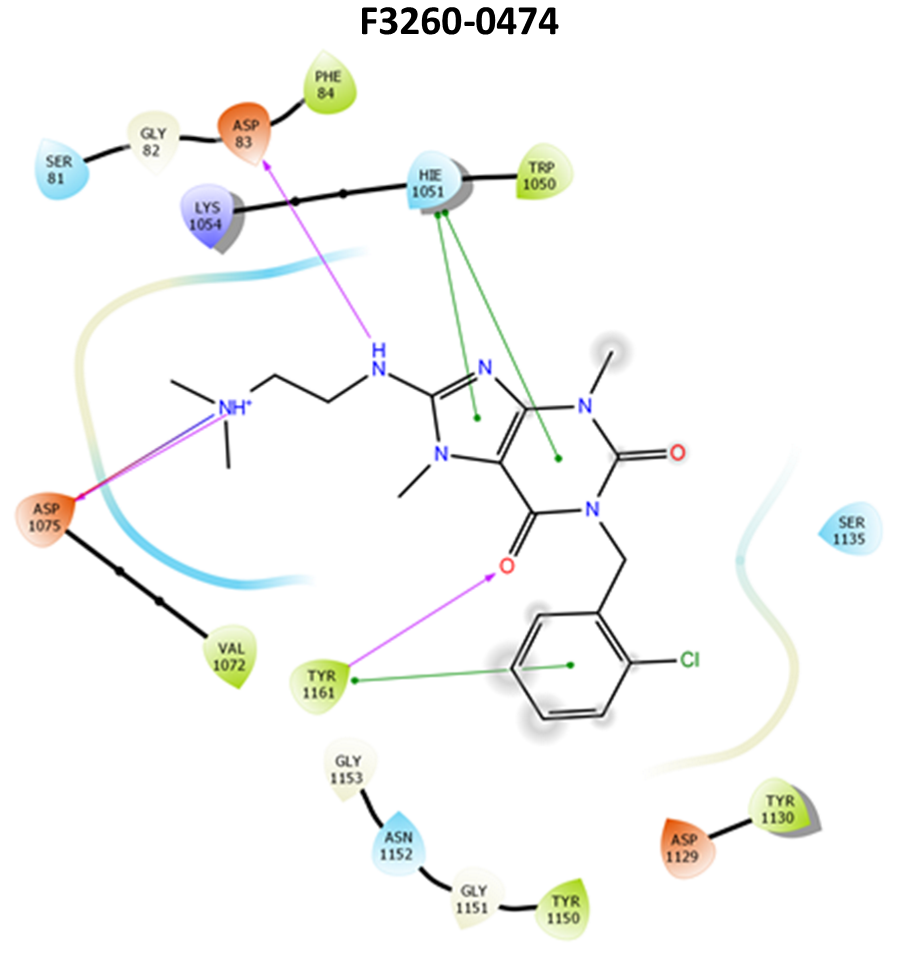

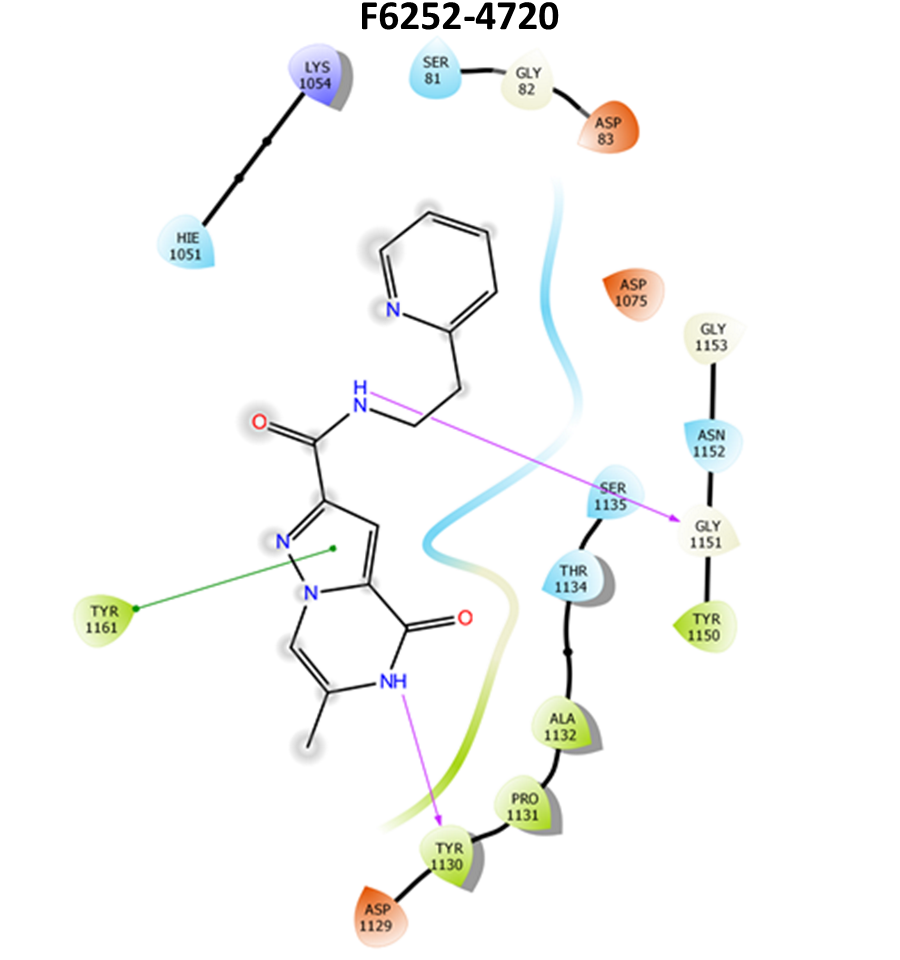

**Figure S2. Molecular interaction of F2477-0054, F3260-0474, and F6252-4720 at the active site of NS2B-NS3 protease.** 2D molecular interaction diagram of NS2B-NS3 protease complexed with F2477-0054, F3260-0474, and F6252-4720. Each compound at the center interacts with the amino acid of NS2B-NS3 protease by H-bond (magenta arrow), salt bridge (red-blue solid line), π-π bond (solid green line), and hydrophobic interactions. F2477-0054 (336.39 g/mol) stabilized at the NS2B-NS3 protease active site by two H-bonds (Asp83 of NS2B and Tyr130 of NS3 protease), two π-π interactions (Tyr161 of NS3 protease), and hydrophobic interactions (Phe84 from NS2B; Val155, Val154, Tyr150, Tyr161, Ala132, Pro131, and Tyr130 from NS3 protease). F3260-0474 (390.87g/mol) is stabilized at the active site by interacting with amino acid residues of NS2B and NS3 protease. It forms three H-bonds (Asp83 of NS2B; Asp75 and Tyr161 of NS3 protease), three π-π interactions (Tyr 161and His51 of NS3 protease, one salt bridge (Asp 75 of NS3 protease), and hydrophobic interactions (Phe84 of NS2B; Val72, Trp50, Tyr161, Tyr150 and Tyr130 of NS3 protease). This compound interacts with two catalytic residues (Asp75 and His51) of NS3 protease. F6252-4720 (297.31 g/mol) is stabilized at the NS2B-NS3 protease active site by two H-bonds (Tyr130 and Gly151 of NS3 protease), one π-π interaction (Tyr161 of NS3 protease), and hydrophobic interactions (Tyr150, Ala132, Pro131, Tyr130 and Tyr161 of NS3 protease).

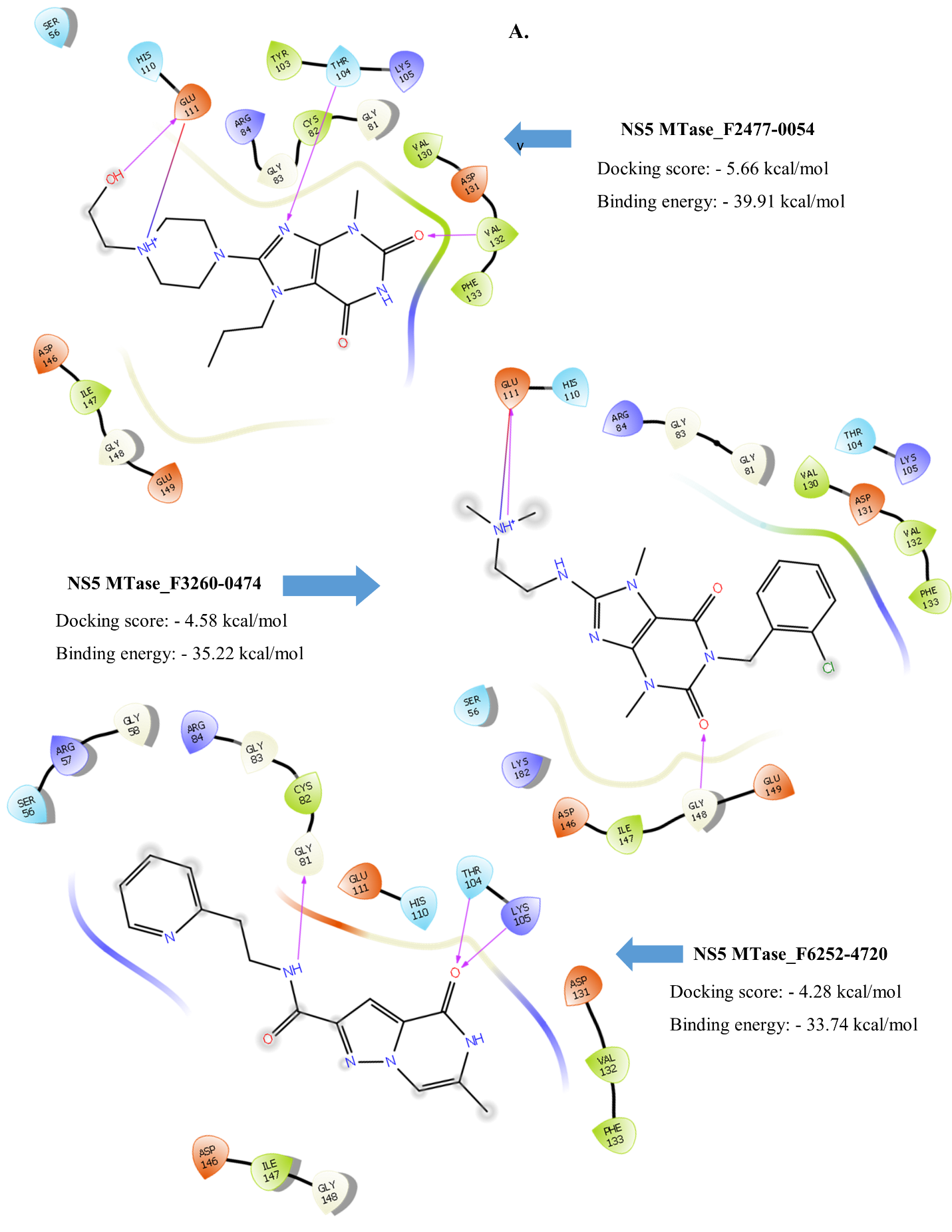

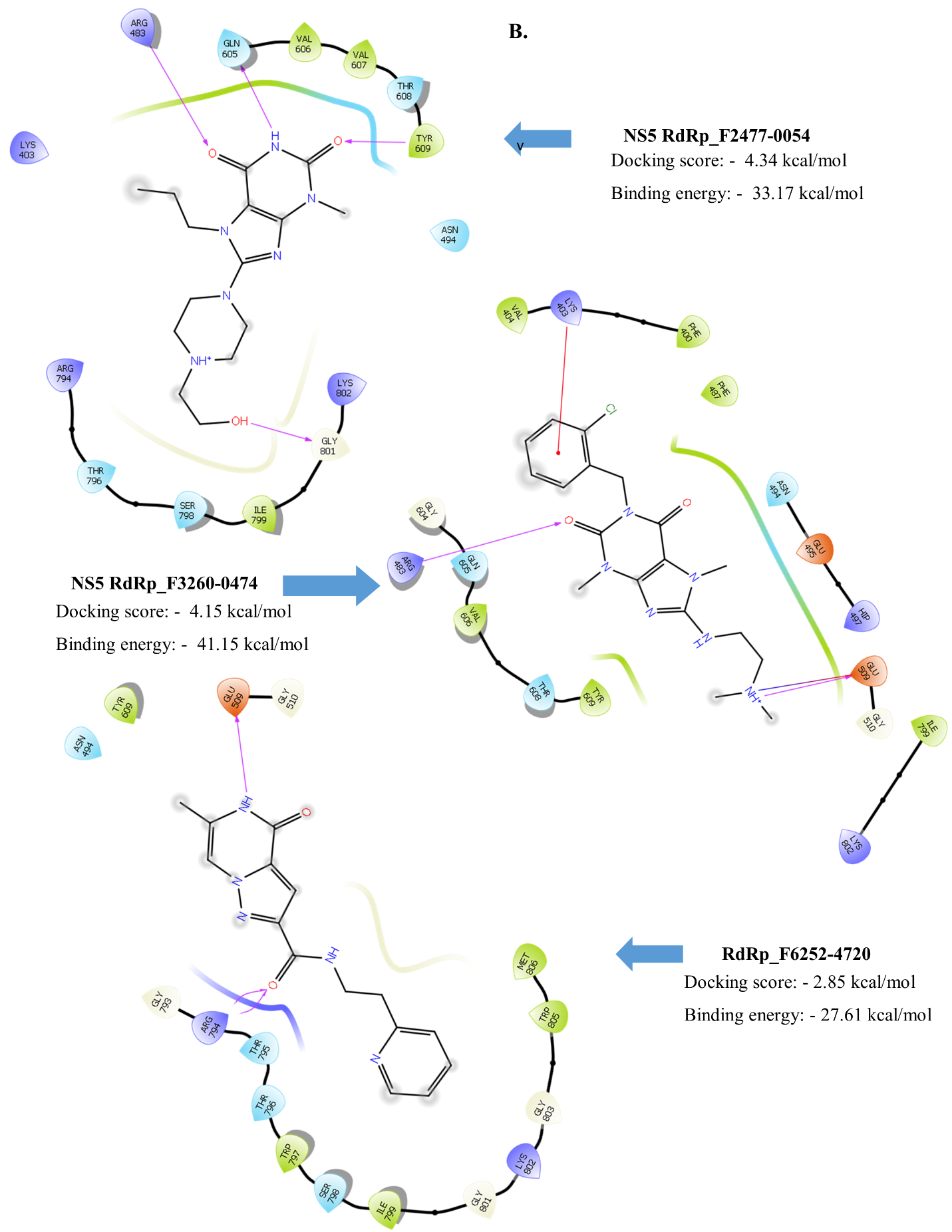

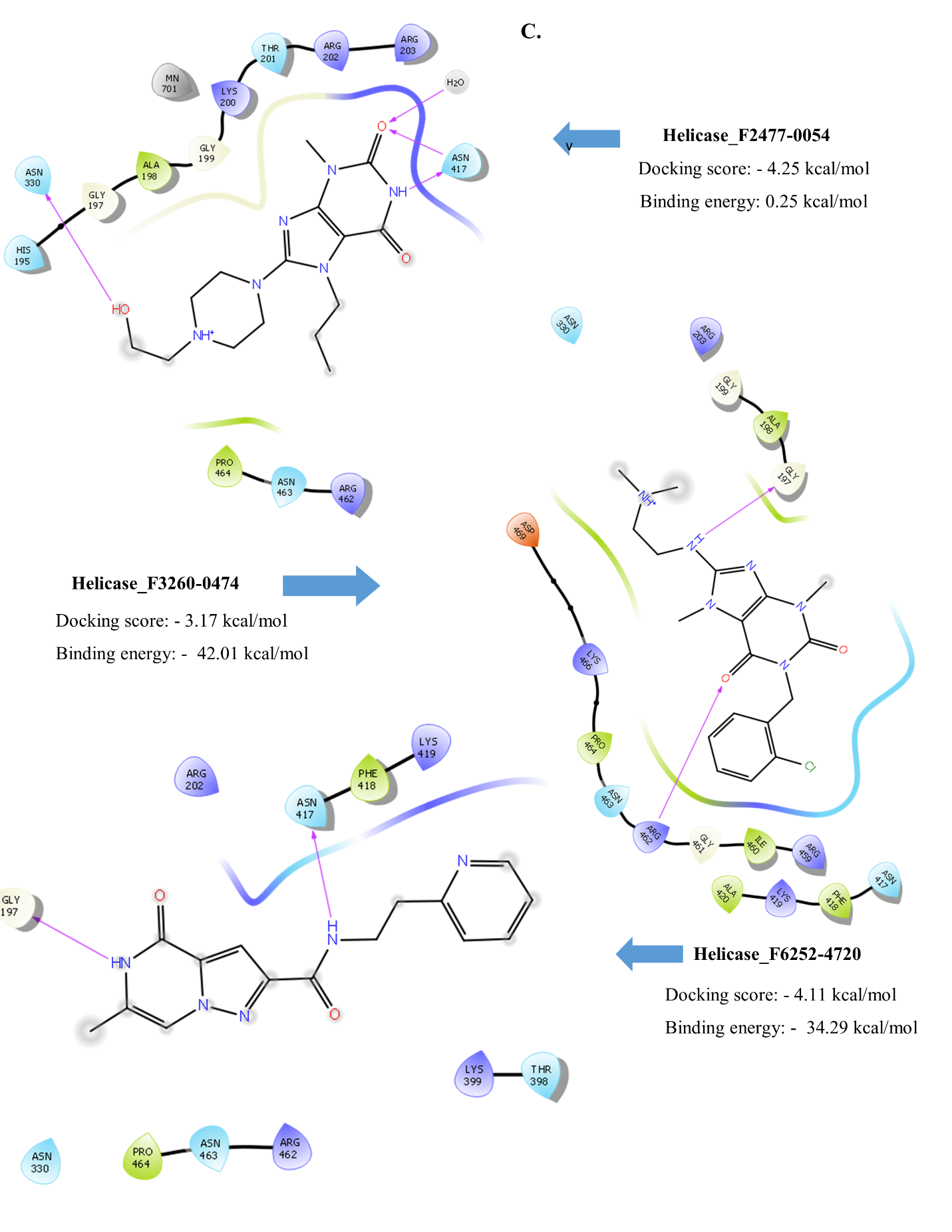
**Figure S3. 2D Molecular interaction of F2477-0054 (A), F3260-0474 (B), and F6252-4720 (C) with Helicase, MTase & RdRp enzyme of ZIKV, respectively.** The compound at the center interacts with the amino acids of these enzymes by H-bond (magenta arrow), salt bridge (red-blue solid line), π-π bond (green solid line), and hydrophobic interactions. (PDB ID: 5KQR for NS5 MTase; 5WZ3 for NS5 RdRp; 5GJC for NS3 helicase).

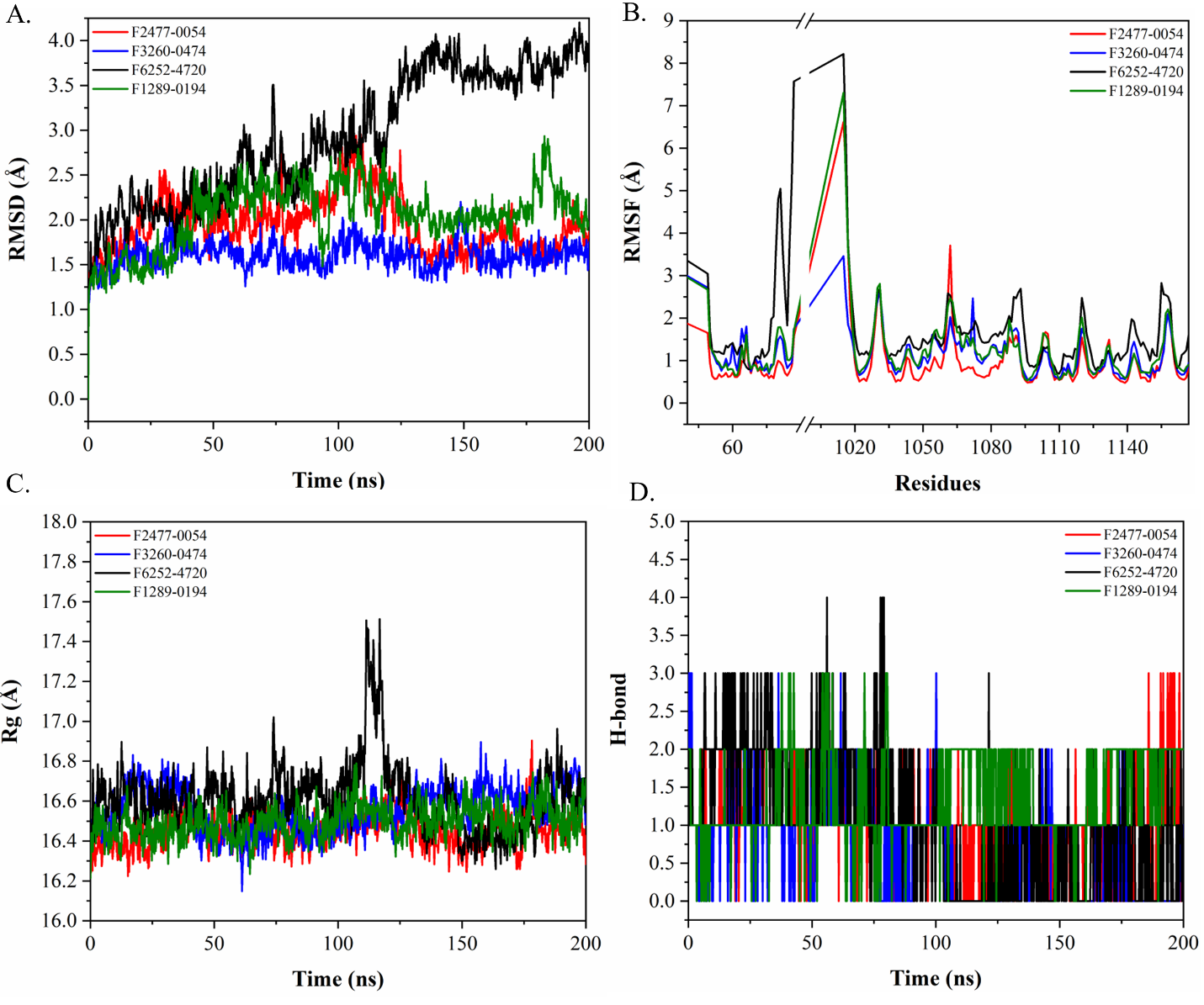
**Figure S4. Molecular dynamics simulation of NS2B-NS3 protease complexed with F1289-0194, F2477-0054, F3260-0474 & F6252-4720**. (A.) RMSD represents the stability of the compounds at the active site. (B.) RMSF of amino acid residue upon addition of compounds. (C.) Rg represents the compaction of NS2B-NS3 protease complexed with the compounds. (D.) represents the loss and gain of the H-bond during simulation. [ns: nanoseconds; in panel B, NS2B residues correspond to 49-87; NS3 protease residues 15-167 are represented as corresponding to 1015-1167].

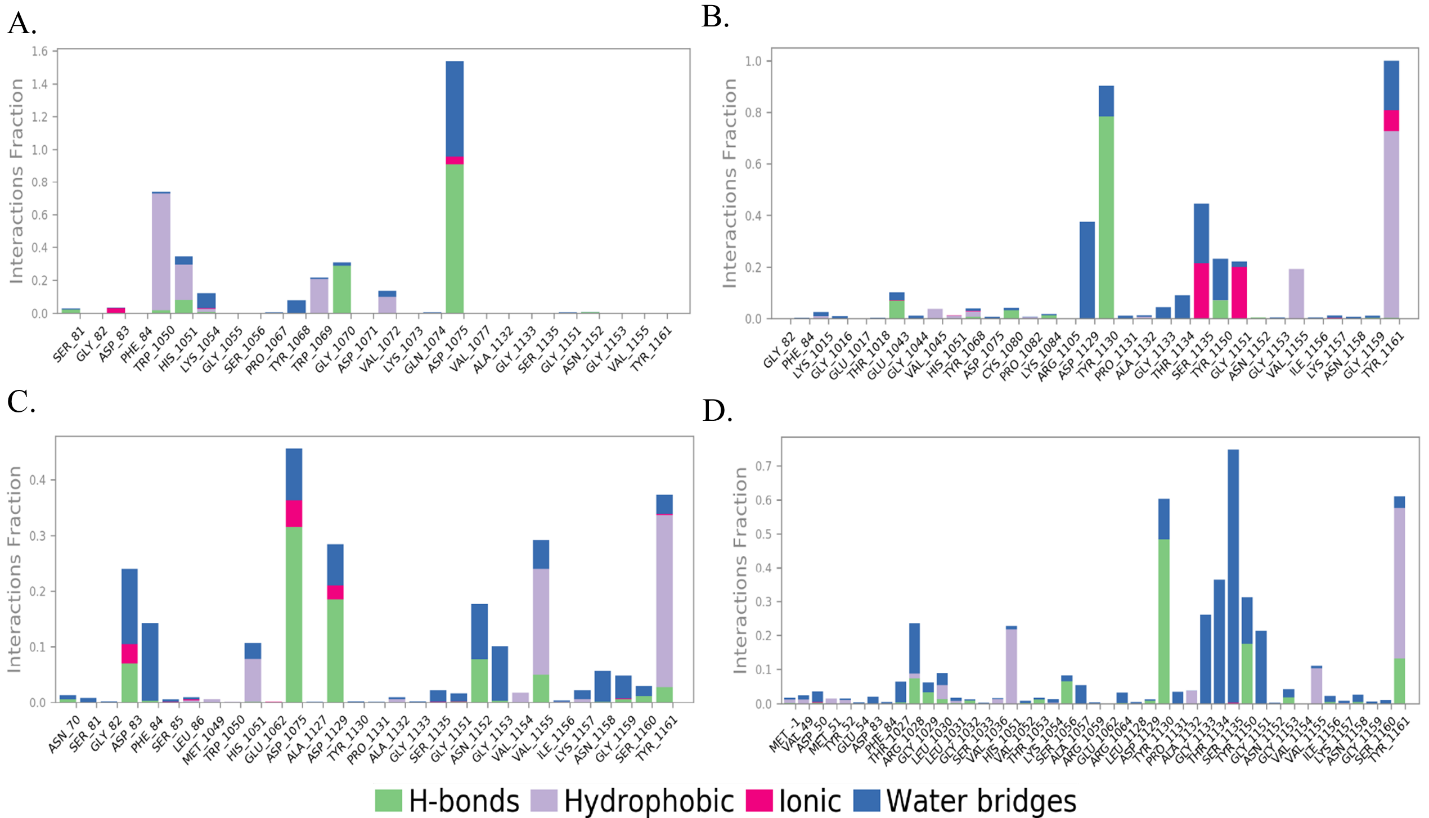

**Figure S5. A bar chart of the interaction of small molecules with the NS2B-NS3 protease residues throughout the simulation of 200 ns.** Panel (A.), (B), (C), & (D) represents the interaction of F1289-0194, F2477-0054, F3260-0474 & F6252-4720, respectively. The interactions like H-bonds, Hydrophobic, ionic, and water bridges are maintained with a different interactions fraction of simulation time. For example, a value of interactions fraction of 1 suggests that the contact is maintained 100 % of the simulation time between the small molecule and NS2B-NS3 protease active site. If the value is more than 1, it indicates that the residue is involved in forming multiple interactions of the same type. [NS2B residues correspond to 49-87; NS3 protease residues 15-167 are represented as corresponding to 1015-1167].

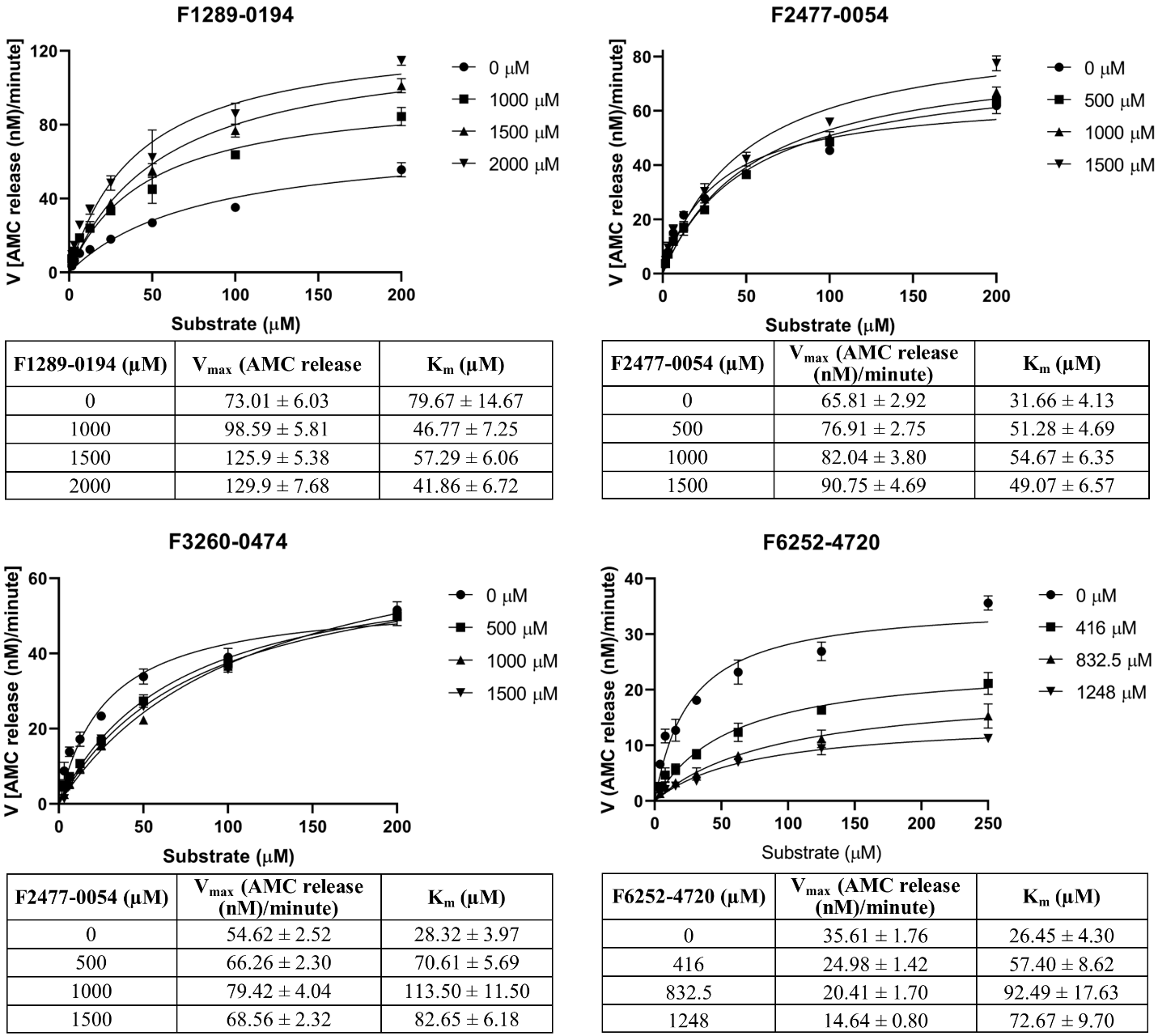

**Figure S6. Enzyme kinetic parameter of NS2B-NS3 protease in the presence of F1289-0194, F2477-0054, F3260-0474 & F6252-4720**. All graphs in the figure were plotted in GraphPad Prism software. The data point in each plot was fitted (R^2^ > 0.90) by the Michaelis-Menten equation to compute the kinetic parameter (K_m_ & V_max_) at the indicated compound concentration. The kinetic parameter's value is represented as ± standard error.

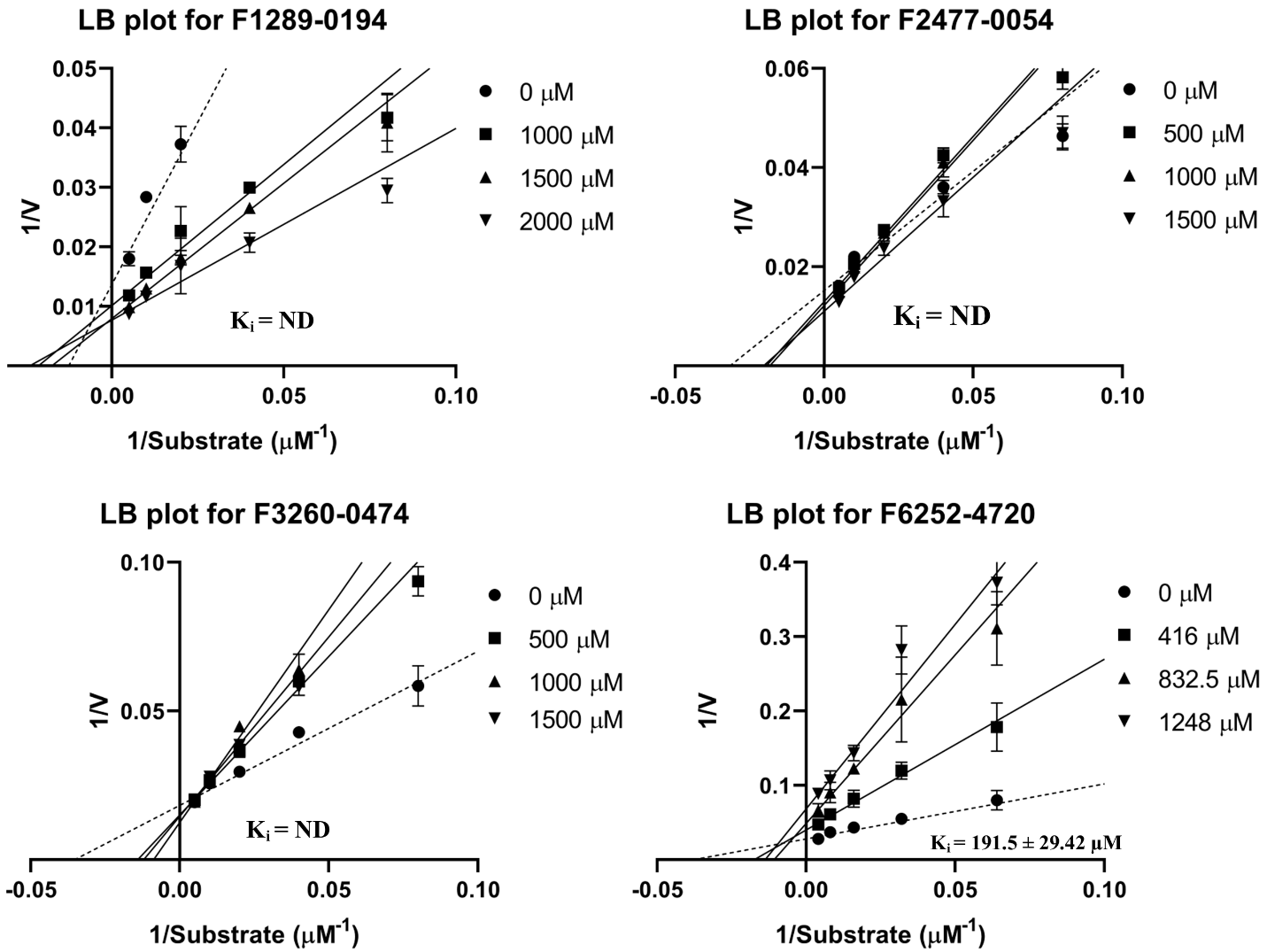

**Figure S7. Lineweaver-Burk (LB) plot for F1289-0194, F2477-0054, F3260-0474 & F6252-4720**. LB plot was prepared from the data point indicated in Figure S6. Based on the LB plot, the inhibition constant (K_i_) was determined for a mixed enzyme inhibition model. All graphs in the figure were plotted in GraphPad Prism software, and the values were represented with ± standard error. [ND = not determined].

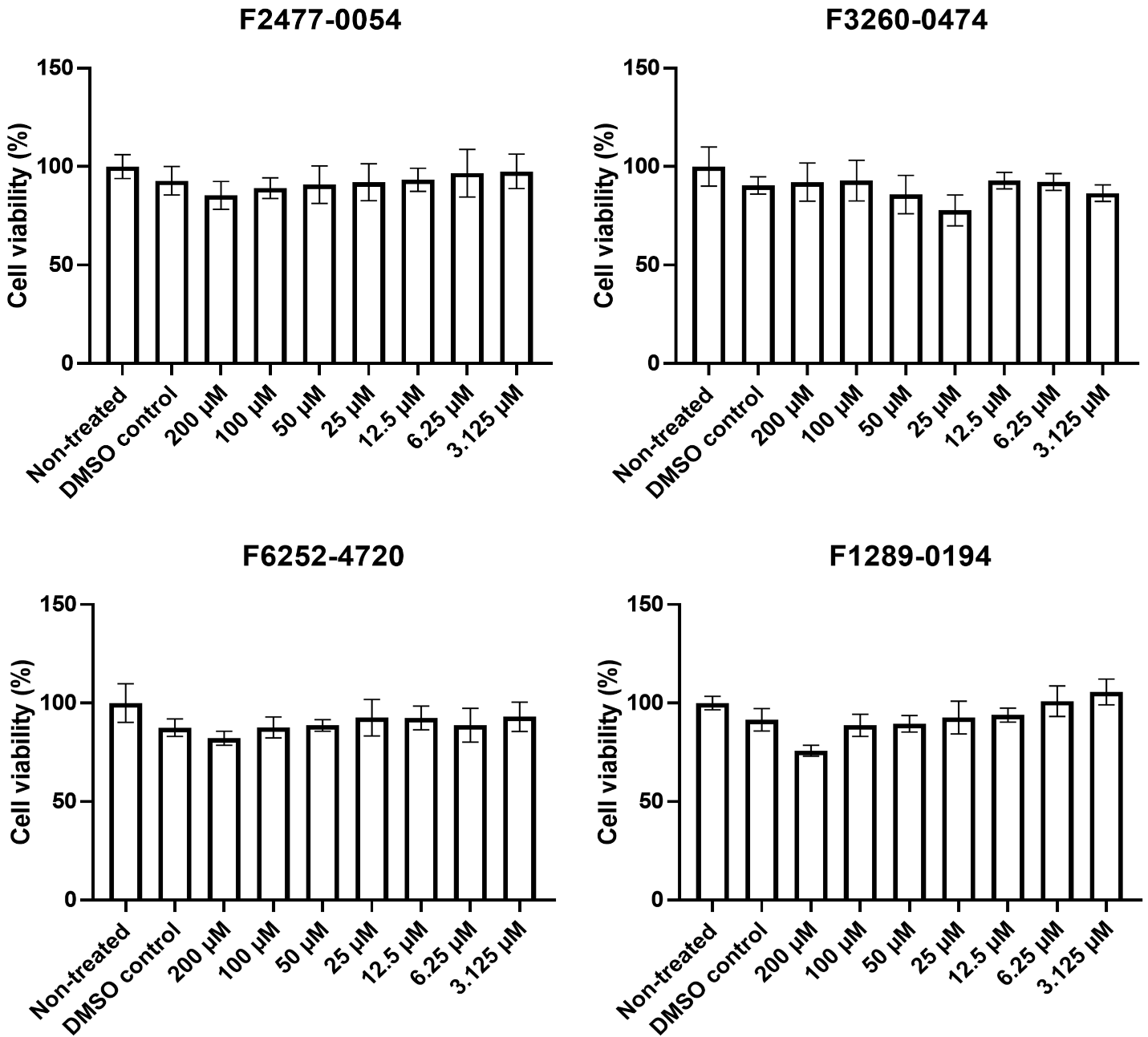
**Figure S8. Cell viability of top hit compounds in Vero cells.** The Vero cells were treated with these compounds (F1289-0194, F2477-0054, F3260-0474, and F6252-4720) for 48 hrs and the MTT assay was performed to estimate cell viability in each case. The percent of cell viability at indicated treatment with respect to the DMSO control was estimated using equation, %Cell viability = [Abs _treated_/ Abs _non-treated_]*100. Experiments were set up in triplicates for two biological replicates. The bar represents the mean ± standard error of the mean.

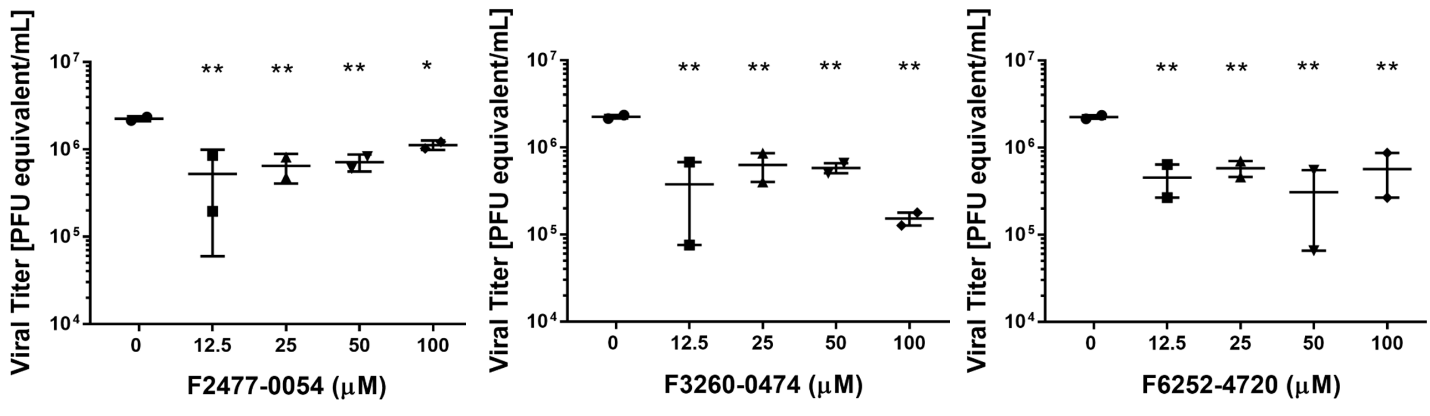
**Figure S9. F2477-0054, F3260-0474, and F6252-4720 inhibit ZIKV replication in Vero cells.** The viral burden in the ZIKV-infected Vero cells after indicated treatment with the compounds F2477-0054, F3260-0474, and F6252-4720 for 48 hpi. RNA from the supernatant of indicated treatment of ZIKV-infected cells was quantified by one-step q-RT-PCR and represented as PFU equivalent/ml. Symbols represent each biological replicate, and the bar represents the mean ± standard error of the mean (n=2; ns: non-significant, ***P < 0.01;* one-way ANOVA Dunnett’s test).

**Table S1**. Docking results of the top 10 poses of the compounds from protease targeted, serine protease targeted, and antiviral combined ligand library and its interaction with amino acid residues of ZIKV NS2B-NS3 protease. Rows indicated in bold font are shortlisted for further investigation using MD simulation and viral assay. [asterisk (*) represents the amino acid residues of NS2B]

| **Serine protease targeted molecules** | | | | | | | | | |
| --- | --- | --- | --- | --- | --- | --- | --- | --- | --- |
| Compound ID | MW (g/mol) | docking score (kcal/mol) | MMGBSA dG Bind (kcal/mol) | H-bond | π-π bond | π-cation | Salt bridge | Hydrophobic interaction | No. of similar scaffolds in the PubChem database |
| F3260-0006 | 472.58 | -7.782 | -41.275 | Hie51, Asp83*, GLY 151 | Tyr161 | - | Asp75, Asp83* | Tyr130, Tyr161, Tyr150, Val154, Val155, Phe84*, Val72, Trp50 | 1859 |
| **F1289-0194** | **308.38** | **-6.979** | **-35.5** | **Asp75, Asp83*, Tyr161, Gly153** | **-** | **-** | **Asp75** | **Trp50, Val72 Tyr161, Val154, Phe84*,** | **6962** |
| F3144-0096 | 490.55 | -6.952 | -45.845 | Ser135, Tyr130 | Tyr161 | - | - | Trp50, Val52, Val72, Tyr150, Tyr161, Val155, Tyr130, Pro131, Ala132 | 2115 |
| **F2477-0054** | **336.39** | **-6.949** | **-27.482** | **Asp83*, Tyr130** | **Tyr161, Tyr161** | **-** | **-** | **Phe84*, Val155, Val154, Tyr150, Tyr161, Ala132, Pro131, Tyr130** | **1805** |
| **F3260-0474** | **390.87** | **-6.85** | **-32.424** | **Asp83*, Asp75, Tyr161** | **Tyr161, Hie51, Hie51** | **-** | **Asp75** | **Val72, Phe84*, Trp50, Tyr161, Tyr150, Tyr130** | **4601** |
| F3260-0126 | 310.35 | -6.809 | -29.499 | Asp75, Asp83*, Tyr161, Gly153 | - | - | Asp75 | Trp50, Val72, Val154, Phe84*, Tyr161 | 2078 |
| F3144-0016 | 324.38 | -6.711 | -30.699 | Asp75, Asp83*, Val155, Gly151 | - | - | Asp75 | Phe84*, Val155, Val154, Tyr161, Val72, Trp50 | 2078 |
| F3260-0001 | 442.51 | -6.697 | -31.684 | Asp83*, Gly151, | Hie51, Tyr161 | - | Asp83*, Asp75 | Trp50, Phe84*, Tyr150, Ala132, Pro131, Tyr130, Tyr161 | 1677 |
| F1687-0119 | 456.54 | -6.643 | -37.671 | Asp75, Hie51, Gly151 | Tyr161 | - | Asp75, asp83 | Phe84*, Val72, Trp50, Tyr150, Val154, Val155, Tyr161, Tyr130, Ala132 | 2073 |
| F2636-0544 | 425.48 | -6.623 | -32.703 | Tyr130, Asp83* | Tyr161, Tyr161 | Hie51 |  | Val155, Tyr150, Val52, Val36, Tyr130, Pro131, Ala132 | 8126 |
| **Protease targeted molecules** | | | | | | | | | |
| F1021-0527 | 284.35 | -7.025 | -44.169 | Phe84*, Gly151, Gly153, Gly153, Tyr161 | Tyr161, Tyr161 | - | - | Tyr130, Pro131, Tyr161, tyr150, Phe84*, Val154, Val155 | 2324 |
| F3176-0232 | 456.54 | -7.003 | -0.798 | Gly153, Tyr161 | Tyr161 | - | Asp75, Asp83* | Trp50, Phe84, Val72, Val154, tyr161, Tyr150, Ala132, Pro131, Tyr130 | - |
| F1021-0527 | 284.35 | -6.905 | -41.182 | Asp83*, Asn152, Gly151 | Tyr161, Tyr161 | - | - | Ala132, Pro131, Tyr130, tyr161, Tyr150, Phe84, val155 | - |
| F3064-0186 | 430.5 | -6.846 | -40.818 | Asp83*, Val155, Gly151, Hie51 | - | - | Asp75 | Ala132, Pro131, Tyr130, tyr161, Tyr150, Phe84, val155, val154, Trp50, Val72 | 2892 |
| F3064-0186 | 430.5 | -6.71 | -31.086 | Asp83*, Asn152, Gly151 | - | - | Asp83*, Asp75 | Ala132, Pro131, Tyr130, tyr161, Tyr150, Phe84, val155, val154, Trp50, Val72 | - |
| F3064-0186 | 430.5 | -6.583 | -38.748 | Asp75, Asp83*, Tyr161 | Tyr161 | - | - | Ala132, Pro131, Tyr130, tyr161, Tyr150, val155, val154, Trp50, Val72, Phe84 | - |
| F3064-0186 | 430.5 | -6.562 | -32.342 | Asp83*, Asp75, Asn152, Gly151, Tyr161 | Tyr161 | - | Asp83* | Tyr130, tyr161, Tyr150, val155, val154, Trp50, Val72, Phe84 | - |
| F3176-0232 | 456.54 | -6.385 | -56.114 | Gly151 | - | - | Asp83*, Asp75 | Ala132, Pro131, Tyr130, tyr161, Tyr150, Trp50, Val72, Phe84 | - |
| F1021-0527 | 284.35 | -6.357 | -32.424 | Asp83*, Asn152, Gly153, Tyr161 | Tyr161, Tyr161 | - | - | Ala132, Pro131, Tyr130, tyr161, Tyr150, val155, Phe84 | - |
| F1021-0527 | 284.35 | -6.352 | -45.31 | Phe84, Gly153, Gly151 | Tyr161, Tyr161 | - | - | Pro131, Tyr130, tyr161, Tyr150, val155, val154, Phe84 | - |
| **Antiviral Combined ligand library** | | | | | | | | | |
| F6524-4573 | 326.31 | -6.801 | -41.065 | Tyr161, Asn152, Gly153, Tyr130 | Tyr161 | - | - | Phe84, Trp50, Tyr150, Val72, Ala132, Pro131, Tyr130, tyr161, | 41 |
| F6492-0928 | 341.45 | -6.453 | -60.45 | Tyr161, Asp83 | Tyr161 | - | Asp75 | Trp50, Tyr150, val155, Val72, Ala132, Pro131, Tyr130, | 858 |
| F6492-0511 | 302.37 | -6.117 | -37.092 | Gly151, Asp83*, Asp75 | Tyr161 | - | Asp75 | Trp50, Tyr150, Val72, Ala132, Pro131, Tyr130, Tyr161, | 9 |
| F6492-0632 | 259.3 | -6.091 | -43.139 | Gly151, Asp83*, Asp75 | Tyr161 | - | Asp75 | Tyr150, Val72, Ala132, Tyr130, Tyr161, | 32 |
| **F6252-4720** | **297.31** | **-6.009** | **-21.483** | **Tyr130, Gly151,** | **Tyr161** | **-** | **-** | **Tyr150, Ala132, Pro131, Tyr130, Tyr161,** | **1367** |
| F6064-1441 | 378.38 | -5.961 | -30.297 | Hie51, Asp83, trp50 | Tyr161 | - | - | Trp50, Tyr150, Val72, Ala132, Pro131, Tyr130, Tyr161, | 18 |
| F6178-4212 | 338.45 | -5.847 | -48.734 | Asn152, Asn152, Tyr130 | - | Tyr161 | - | Trp50, Tyr150, Val72, Ala132, Pro131, Tyr130, tyr161, | 685 |
| F6412-6605 | 329.39 | -5.843 | -35.312 | Asp129, Tyr130 | Tyr161 | - | Asp75, asp83* | Phe84, Trp50, Tyr150, Val72, Val154, Ala132, Pro131, Tyr130, tyr161, | 3106 |
| F6506-1248 | 321.35 | -5.843 | -34.838 | Gly151, Tyr161, Tyr130 | - | - | - | Phe84, Trp50, Val154, Val155, Ala132, Pro131, Tyr130, tyr161, | 67 |
| F6472-8364 | 355.86 | -5.838 | -40.272 | Hie51, Tyr161, | Tyr161 | - | Asp75 | Trp50, Tyr150, Val72, Pro131, Tyr130, tyr161, | 96 |
